# Spatiotemporal transcriptomic profiling and modeling of mouse brain at single-cell resolution reveals cell proximity effects of aging and rejuvenation

**DOI:** 10.1101/2024.07.16.603809

**Authors:** Eric D. Sun, Olivia Y. Zhou, Max Hauptschein, Nimrod Rappoport, Lucy Xu, Paloma Navarro Negredo, Ling Liu, Thomas A. Rando, James Zou, Anne Brunet

## Abstract

Old age is associated with a decline in cognitive function and an increase in neurodegenerative disease risk^1^. Brain aging is complex and accompanied by many cellular changes^2–20^. However, the influence that aged cells have on neighboring cells and how this contributes to tissue decline is unknown. More generally, the tools to systematically address this question in aging tissues have not yet been developed. Here, we generate spatiotemporal data at single-cell resolution for the mouse brain across lifespan, and we develop the first machine learning models based on spatial transcriptomics (‘spatial aging clocks’) to reveal cell proximity effects during brain aging and rejuvenation. We collect a single-cell spatial transcriptomics brain atlas of 4.2 million cells from 20 distinct ages and across two rejuvenating interventions—exercise and partial reprogramming. We identify spatial and cell type-specific transcriptomic fingerprints of aging, rejuvenation, and disease, including for rare cell types. Using spatial aging clocks and deep learning models, we find that T cells, which infiltrate the brain with age, have a striking pro-aging proximity effect on neighboring cells. Surprisingly, neural stem cells have a strong pro-rejuvenating effect on neighboring cells. By developing computational tools to identify mediators of these proximity effects, we find that pro-aging T cells trigger a local inflammatory response likely via interferon-γ whereas pro-rejuvenating neural stem cells impact the metabolism of neighboring cells possibly via growth factors (e.g. vascular endothelial growth factor) and extracellular vesicles, and we experimentally validate some of these predictions. These results suggest that rare cells can have a drastic influence on their neighbors and could be targeted to counter tissue aging. We anticipate that these spatial aging clocks will not only allow scalable assessment of the efficacy of interventions for aging and disease but also represent a new tool for studying cell-cell interactions in many spatial contexts.

## Introduction

Brain aging is associated with a marked increase in the risk of neurodegenerative diseases, including Alzheimer’s disease and other forms of dementia^1^. Previous studies have interrogated the molecular changes that occur during brain aging at single-cell resolution^2–20^. These datasets provide rich information on age-related cellular changes in the brain, but they lack insight into the spatial context–especially at scale. Thus, we are missing a systematic understanding of spatiotemporal changes in the brain during aging, including changes in local cell neighborhoods and cell-cell interactions. Such interactions are particularly important for brain aging and disease given the complex organization of this organ, the large number of different cell types, and the potential for rare cells having a strong influence on their neighbors.

The advent of high-throughput spatial omics holds great promise for characterizing spatial interactions^21^. While recent studies on spatial brain aging have helped clarify cellular and regional changes with age^22–24^, they have provided either spatial single-cell resolution^22^ or temporal resolution^23,24^, but not both. The lack of high spatiotemporal resolution at single-cell level in current studies makes it difficult to understand the full range of cell type-specific changes and interactions that occur throughout life, particularly in geriatric ages where cognitive decline and the onset of neurodegenerative disease are most prominent. Importantly, advanced computational tools to analyze the wealth of data from spatial omics and capture spatial interactions during aging are missing and this has hindered new discoveries. Here we generate the first spatially resolved single-cell transcriptomics atlas of the mouse brain across adult life and in response to rejuvenating interventions (exercise and partial reprogramming), and we develop new computational tools to discover prominent cell proximity effects in spatial datasets.

## Results

### Spatiotemporal single-cell transcriptomic profiling of the aging mouse brain

We generated a single-cell spatial transcriptomics atlas of the aging mouse brain across the entire lifespan (Fig. 1a). We collected coronal brain sections from male mice at 20 different ages tiling the entire lifespan (two independent cohorts of mice, see Methods), as well as sagittal brain sections from male mice at 6 different ages. We obtained coronal sections that contained the cortex (CTX), striatum and adjacent regions (STR), corpus callosum and anterior commissure (CC/ACO), and lateral ventricles (VEN). We obtained sagittal sections that contained not only the aforementioned regions but also additional regions (olfactory bulb, rostral migratory stream, brain stem, and cerebellum). To generate spatial transcriptomic data at single-cell resolution, we used the MERFISH technology^25–27^ and measured transcripts for 300 genes across entire coronal or sagittal sections. This MERFISH 300 gene panel was designed to contain markers for different cell types or subtypes, genes in aging-related pathways, genes in important processes not previously linked to aging, and genes identified by analysis of single-cell RNA-seq atlases of murine brain aging (see Methods for selection of genes and Supplementary Table 1 for the full list of genes and the rationale for their choice).

**Figure 1:**
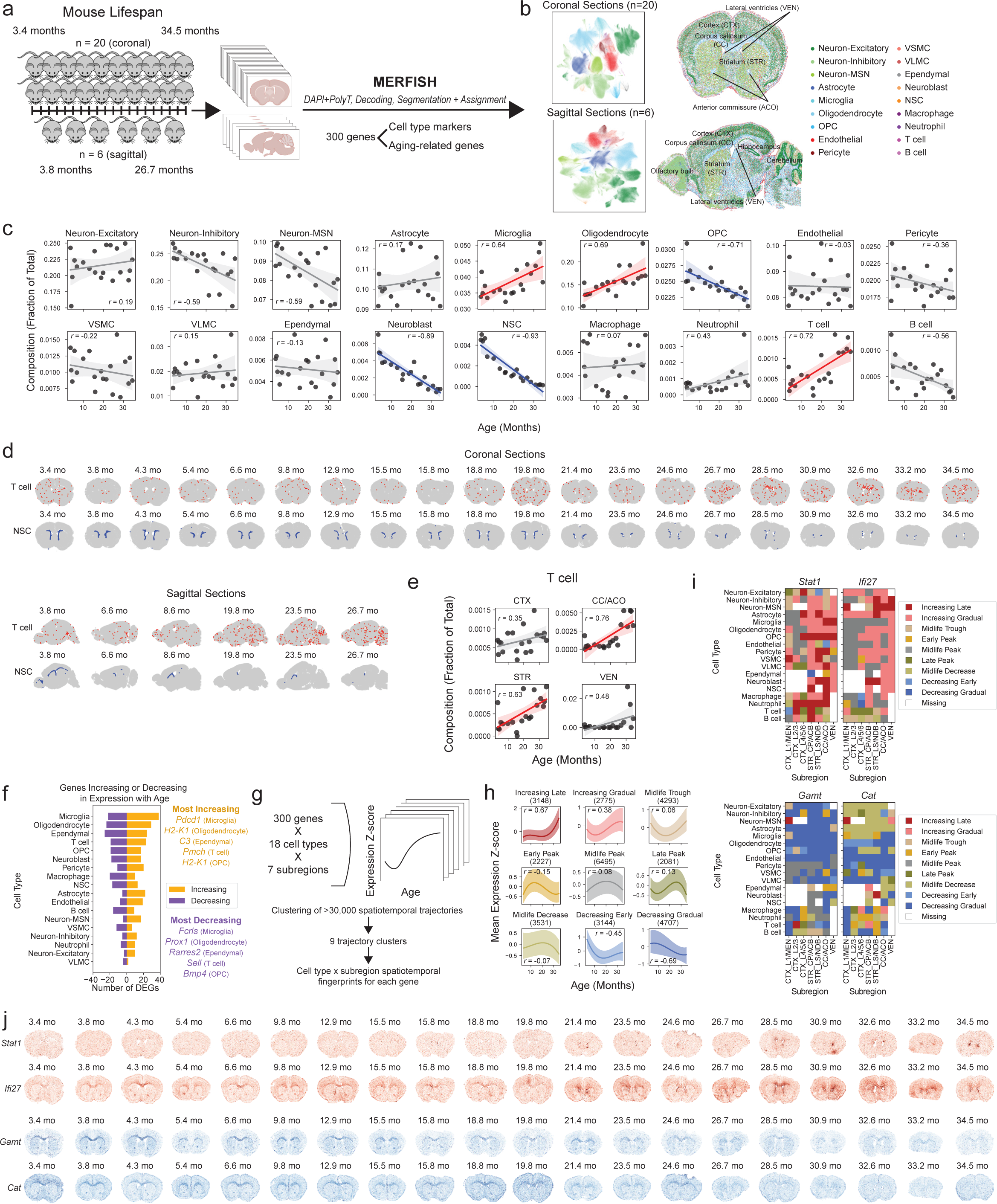
Spatially resolved single-cell transcriptomic profiling of the brain across lifespan. **a,** Experimental workflow for generating spatial transcriptomics data from mouse brains collected across the adult lifespan using MERFISH. Twenty male mice from 3.4 to 34.5 months for coronal sections (in two independent cohorts). Six male mice from 3.8 to 26.7 months for sagittal sections. **b,** Visualization of single cells colored by the identified cell type under two-dimensional UMAP coordinates across all coronal section samples (top left) and across all sagittal section samples (bottom left), and visualizations under spatial coordinates for one representative coronal section (top right) with region labels and one representative sagittal section (bottom right) with some anatomic structures annotated. See Extended Data Fig. 1e,f for additional visualizations by cell type and Extended Data Fig. 3a,b for visualizations by region and subregion. **c,** Global cell type composition changes in the coronal section dataset with each dot representing an individual mouse. Line and shaded region correspond to regression line of best fit and corresponding 95% confidence interval. Pearson correlations between cell type composition and age are reported. Significant increases in cell type proportion with age are in red and significant decreases in cell type proportion with age are in blue (Bonferroni corrected *P*-value < 0.05). **d,** Scatter plot of cells by their spatial coordinates across all ages for all coronal sections (top) and all sagittal sections (bottom) with cells colored by cell type: T cell (red), neural stem cell (NSC) (blue), other cell type (gray). **e,** Region-specific composition changes in the coronal section dataset shown for T cells across the four regions (cortex, striatum and adjacent regions, corpus callosum and anterior commissure, lateral ventricles) with the same plotting and statistical parameters as in (**c**). **f,** Number of genes increasing or decreasing with age for each cell type (coronal sections) determined by the Spearman correlation of age and log-normalized gene expression. “Increasing” corresponds to genes with Spearman correlation greater than 0.3 and strictly positive 95% confidence interval, and “Decreasing” corresponds to genes with Spearman correlation less than -0.3 and strictly negative 95% confidence interval. Select genes with the largest positive and largest negative Spearman correlation coefficients are shown for the top five cell types. **g,** Schematic illustration of the clustering approach for identifying spatiotemporal gene expression trajectories defined by distinct combinations of gene, cell type, and subregion (subregions as defined in Extended Data Fig. 3b). **h,** Median of the mean gene expression z-scores (see Methods) across age of all gene, cell type, and subregion combinations (coronal sections) split into nine annotated clusters (determined by k-means clustering). See Extended Data Fig. 4d,e for optimization of number of clusters. Shaded regions correspond to interquartile range in expression. All trajectories were smoothed using interpolating B-splines. Pearson correlation (*r*) between median gene expression z-score and age is shown, and the number of trajectories within each cluster is noted inside parentheses. **i,** Heatmaps showing trajectory cluster membership (coronal sections) across different cell types and subregions for interferon-genes *Stat1* (top left) and *Ifi27* (top right), which increase in expression with age, and creatine synthesis associated gene *Gamt* (bottom left) and antioxidant enzyme gene *Cat* (bottom right), which decrease in expression with age. Additional genes are provided in Extended Data Fig. 4g. **j,** Scatter plot of cells by their spatial coordinates across all coronal sections and ages with cells colored by scaled log-normalized gene expression of *Stat1* and *Ifi27* (red) and of *Gamt* and *Cat* (blue).

Our spatiotemporal atlas of the aging mouse brain yielded a total of 2.3 million high quality cells from coronal sections across 20 ages and sagittal sections across 6 ages (see Supplementary Tables 2-4). MERFISH measurements were highly reproducible across adjacent coronal sections (Extended Data Fig. 1a) and the log-normalized gene expression quantiles were similar across different sections (Extended Data Fig. 1b).

To identify cell types, we performed Leiden clustering^28^ on the standardized spatial gene expression profiles of individual cells and annotated the resulting clusters using pre-defined marker genes that were included in the MERFISH gene panel (Extended Data Fig. 1c,d, Supplementary Table 5). In the coronal dataset, we identified 18 cell types including neuronal cell types: excitatory neurons (Neuron-Excitatory), inhibitory neurons (Neuron-Inhibitory), medium spiny neurons (Neuron-MSN); glial cell types: astrocytes, oligodendrocytes and oligodendrocyte progenitor cells (OPC); cell types localized to the lateral ventricles, which contain the neurogenic niche: neural stem cells (NSCs), neuroblasts, ependymal cells; cell types involved in brain vasculature: endothelial cells, pericytes, vascular smooth muscle cells (VSMC), vascular leptomeningeal cells (VLMC); and immune cell types: microglia (the resident immune cell of the brain), T cells, B cells, macrophages, and neutrophils (Fig. 1b, Extended Data Fig. 1e). We identified the same cell types except for neutrophils in the sagittal dataset (Fig. 1b). Importantly, our atlas and analysis identified both abundant cell types (e.g. excitatory and inhibitory neurons, oligodendrocytes, astrocytes, microglia) and rare cell types (e.g. T cells, B cells, neutrophils, ependymal cells, neural stem cells, neuroblasts), some of which have not been studied in previous spatial transcriptomics atlases of the brain^22,29–34^. The cell types we identified localized to their expected spatial regions. For example, excitatory neurons were predominantly found in the cortex, while oligodendrocytes were most densely populated in the white matter tracts (Fig. 1b, Extended Data Fig. 1e). The spatial localization of the main cell types identified in our dataset was consistent with that of the same cell types in existing spatial transcriptomics studies^22,29,31^ (Extended Data Fig. 1f). Our immunofluorescence staining and imaging also confirmed the spatiotemporal expression of specific markers in the panel (Extended Data Fig. 2). Together, these results confirm the validity of cell type markers and identification in our spatial transcriptomics dataset.

To determine how cells localize to different regions of the brain, we performed unbiased clustering of cells based on cell neighborhood abundances^22^. This clustering resulted in annotation of seven anatomic subregions that were manually grouped into four regions: 1) white matter tracts of the corpus callosum and anterior commissure (CC/ACO), 2) three subregions of the cortex (cortex layer 1 and meninges [CTX_L1/MEN], cortex layer 2/3 [CTX_L2/3], cortex layer 4/5/6 [CTX_L4/5/6]), 3) two subregions of the striatum and adjacent regions (caudoputamen and nucleus accumbens [STR_CP/ACB] and septal nucleus and diagonal band nucleus [STR_LS/NDB]), and 4) the lateral ventricles (VEN) (Fig. 1b, Extended Data Fig. 3a,b). Known cortical layer and neuronal markers exhibited similar spatial expression patterns between our dataset and the Allen brain *in situ* hybridization atlas^35^ (Extended Data Fig. 3c,d, Supplementary Table 1), confirming the annotation of the cortical layers.

In concordance with previous studies^2,4,5,22,36^, we observed a substantial global increase in the proportion of microglia, oligodendrocytes, and T cells during aging, as well as a substantial global decrease in the proportion of OPCs, NSCs, and neuroblasts in the coronal section dataset (Fig. 1c, Supplementary Table 6). Notably, T cells and NSCs exhibited the strongest changes in proportion with age. T cells were found across all sampled regions and substantially increased in proportion with age (Pearson correlation test *P* = 3.8×10^−4^, *r* = 0.72 with 95% confidence interval [0.40, 0.88]) (Fig. 1c,d). NSCs were generally localized to the lateral ventricles throughout life and substantially decreased in proportion with age (Pearson correlation test *P* = 4.9×10^−9^, *r* = -0.93 with 95% confidence interval [-0.97, -0.82]) (Fig. 1c,d).

We observed region-specific cell type proportion changes with age (Supplementary Table 6). For example, oligodendrocytes primarily increased in the cortex and striatum/adjacent regions during aging (Extended Data Fig. 3e), whereas OPCs primarily decreased in the cortex, corpus callosum/anterior commissure, and striatum/adjacent regions (Extended Data Fig. 3f). Interestingly, both T cells and microglia primarily increased in the corpus callosum/anterior commissure and striatum/adjacent regions with age (Fig. 1e, Extended Data Fig. 3g).

To determine whether similar changes in cell type proportion during aging could be observed in other regions of the brain, we used our sagittal section dataset, which contained regions like the olfactory bulb and cerebellum that were not captured in the coronal section dataset. The proportion of several cell types changed with age in sagittal sections, and these changes were largely consistent with those observed in coronal sections (Extended Data Fig. 3h). T cells were also found to increase in several regions of the brain with age in sagittal sections (Fig. 1d). NSCs were present mostly in the subventricular zone of the lateral ventricles, the rostral and caudal migratory streams, and to a lower extent in the dentate gyrus in sagittal sections (Fig. 1d). Similar to what we observed in coronal sections, NSCs decreased in the subventricular zone within the lateral ventricles in sagittal sections (Fig. 1d).

Together, these datasets represent a spatially resolved single-cell atlas of multiple brain regions across life consisting of around 2.3 million high-quality cell gene expression profiles, including rare cell types such as T cells, B cells, neutrophils, neural stem cells, neuroblasts, and ependymal cells, and revealing striking changes in cell type proportions with age.

### Spatiotemporal gene expression changes during aging

We identified genes that increased or decreased in expression during aging in each cell type using Spearman correlation to account for both linear and nonlinear changes with age (see Methods, Supplementary Table 7). Genes that increased in expression with age were generally involved in immune response (Extended Data Fig. 4a, Supplementary Table 8), while genes that decreased in expression with age were generally involved in cellular metabolism and development pathways (Extended Data Fig. 4b, Supplementary Table 8). We found that microglia, the resident immune cell type of the brain, experienced the largest number of genes that increased or decreased in expression with age of any cell type (Fig. 1f). In microglia, *Pdcd1*, which encodes the PD-1 receptor, was one of the most strongly upregulated genes with age (Spearman correlation with age *ρ* = 0.93), and *Fcrls*, which encodes a Fc receptor-like molecule, was the most strongly downregulated gene with age (Spearman correlation with age *ρ* = -0.76).

The magnitude of transcriptomic changes in different cell types varied across subregions. The corpus callosum and anterior commissure region, which consists of white matter tracts, experienced the largest gene expression changes across multiple cell types with age (Extended Data Fig. 4c), in line with findings of white matter tracts as hotspots of aging in the brain^24^.

To characterize the shape of gene expression changes during aging (hereafter ‘trajectories’), we leveraged the high temporal resolution of our spatial transcriptomics dataset. By clustering gene expression trajectories across all 20 ages for each combination of gene, cell type, and spatial subregion (as defined in Extended Data Fig. 3a), we identified nine different clusters representing archetypes of gene expression trajectories (Fig. 1g,h, Supplementary Fig. 1). The number of clusters was chosen to optimize for distinct gene expression trajectories (Extended Data Fig. 4d,e). These trajectories include genes with expression that increases late in life (“increasing late”), increases gradually throughout life (“increasing gradual”), has the lowest expression in midlife (“midlife trough”), peaks in early life (“early peak”), peaks in midlife (“midlife peak”), peaks in late life (“late peak”), decreases after midlife (“midlife decrease”), decreases early in life (“decreasing early”), and decreases gradually through life (“decreasing gradual”) (Fig. 1h). Within a given cell type and subregion, the genes represented in these spatiotemporal trajectory clusters were associated with distinct biological processes (Extended Data Fig. 4f, Supplementary Table 9). For example, for oligodendrocytes in the corpus callosum/anterior commissure, “decreasing early” genes are implicated in development pathways while “decreasing gradual” genes are involved in stress response and DNA damage repair; and “increasing gradual” genes are implicated in many cell signaling and development pathways while “increasing late” are involved in immune response (Extended Data Fig. 4f, Supplementary Table 9). Some of the changes that occur earlier (e.g. decreased stress response and DNA repair) might in fact be responsible for later changes (e.g. increased immune response).

This analysis provides a spatiotemporal transcriptomic fingerprint for any given gene, showcasing its temporal expression pattern for each combination of subregion and cell type. For example, the spatiotemporal fingerprints for the interferon-response genes *Stat1* and *Ifi27* show both “increasing late” (dark red) and “increasing gradual” (light red) trajectories with age across virtually all cell types and subregions (Fig. 1i). Conversely, the spatiotemporal fingerprint for *Gamt*, a gene involved in creatine synthesis, broadly shows “decreasing gradual” (dark blue) trajectories with age, while the antioxidant enzyme gene *Cat* shows cell type-specific “decreasing gradual” (dark blue) expression in oligodendrocytes, OPCs, and vascular cell types across most subregions (Fig. 1i). Several genes exhibit cell type-specific gene and/or subregion-specific expression trajectories with age (see Extended Data Fig. 4g for examples). For example a gene like *H2-K1* (Extended Data Fig. 4g), which encodes a histocompatibility protein involved in antigen presentation, shows “increasing late” (dark red) expression in specific cell types (e.g. astrocytes, microglia, oligodendrocytes, OPCs) and the timing of expression of this gene occurs at geriatric ages where cognitive decline and the onset of neurodegenerative disease are most prominent. The expression of these example genes can also be visualized directly on brain sections (Fig. 1j), confirming the cell type-specific and region-specific changes in gene expression highlighted by the spatiotemporal fingerprints. Using immunofluorescence staining, we also validated the spatiotemporal expression of STAT1 in young, middle-aged, and old brains (Extended Data Fig. 2). This analysis highlights diverse spatiotemporal trajectories during aging at the gene, cell type, and region level.

Overall, our atlas provides a spatiotemporally resolved view of brain aging across 18 cell types, which enables us to test aspects that could not be investigated previously, including interactions between specific cells during aging.

### Building and evaluating new spatial single-cell aging clocks

There is a dearth of computational tools for the analysis of high-dimensional spatiotemporal transcriptomics data. To address this challenge, we developed new machine learning approaches for spatiotemporal analyses. To quantitatively measure the biological age of each cell in the brain, we built machine learning models trained on spatially preprocessed gene expression data to predict an individual’s age for each cell (hereafter ‘spatial aging clocks’). The MERFISH panel we designed includes several genes with a causal role in aging^37^ (see Methods and Supplementary Table 1), which are helpful for generating more biologically meaningful aging clocks^38,39^.

To preserve spatial information while maximizing the performance of single-cell aging clocks, we developed a soft spatial pseudobulking procedure, referred to as SpatialSmooth. This method involves iterative smoothing of gene expression values along a spatial graph of neighboring cells of the same cell type as a preprocessing procedure before training cell type-specific aging clock models on the resulting smoothed single-cell transcriptomes (Fig. 2a, see Methods for details). Aging clocks developed using the SpatialSmooth method on the coronal section dataset yielded high performance (*R* > 0.7) across 14 of the 18 cell types, including very rare cell types such as T cells, NSCs, and neuroblasts (Fig. 2b). Aging clock performance was generally robust to the choice of the number of nearest neighbors used in SpatialSmooth (Extended Data Fig. 5a) and was similar when trained and evaluated on the two independent cohorts in the coronal section dataset (Extended Data Fig. 5b,c). Aging clocks trained using SpatialSmooth substantially outperformed those trained directly on the spatial single-cell transcriptomes of the coronal section dataset (Fig. 2c). These spatial aging clocks generally outperformed previous cell type-specific transcriptomic aging clocks^3^ (Extended Data Fig. 5d), and could be generated from substantially more cell types, even rare ones (e.g. ependymal cells, macrophages, T cells, and pericytes). For all downstream applications, we used the 14 high-performing spatial aging clocks based on SpatialSmooth.

**Figure 2:**
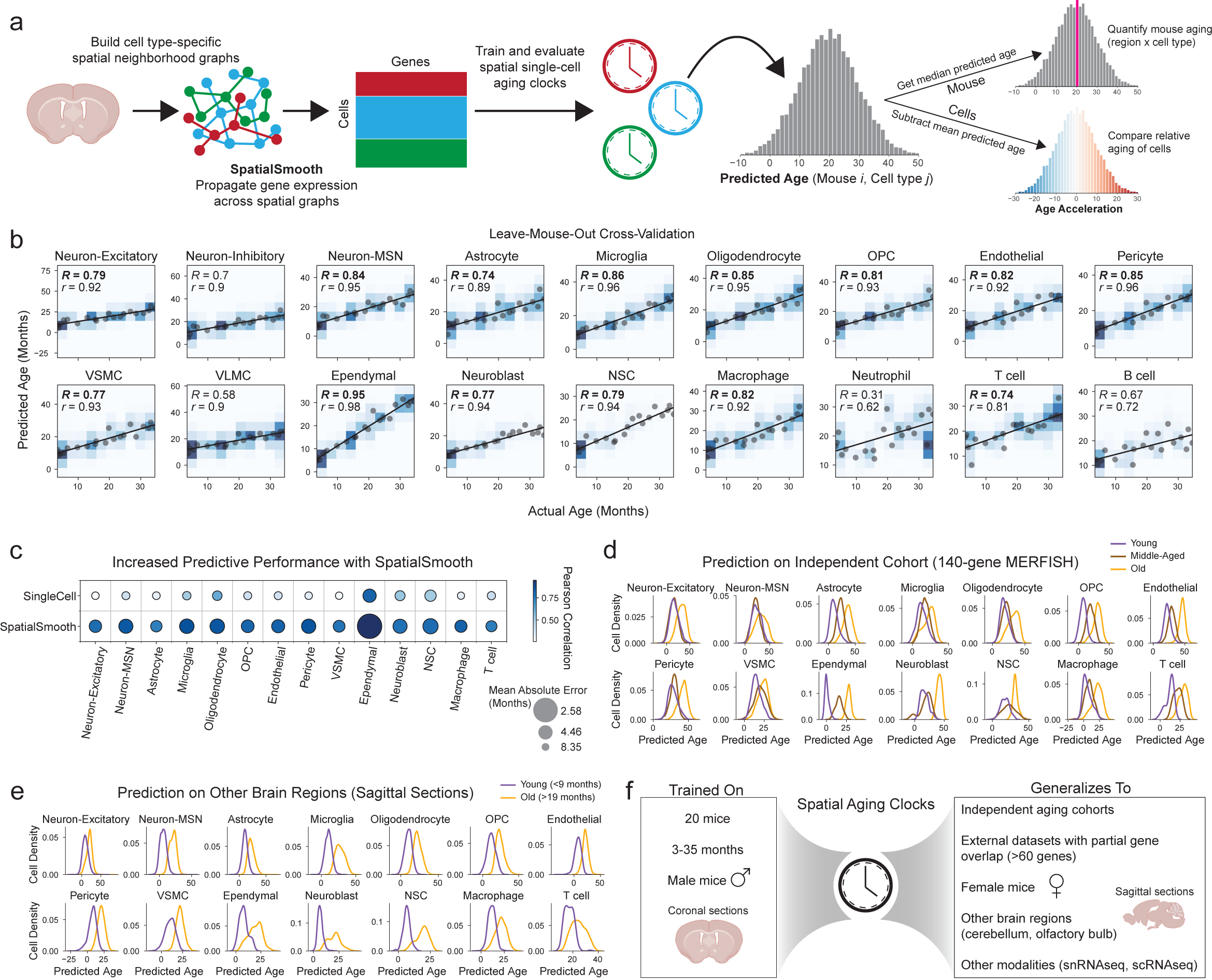
Spatial single-cell transcriptomic aging clocks. **a,** Computational workflow for building spatial single-cell transcriptomic aging clocks from the coronal section dataset, including spatial smoothing of gene expression followed by cell type-specific training of spatial aging clock models. The median predicted age can be used to compare aging between cell types, regions, and different mice. Predicted ages obtained from spatial aging clocks can also be transformed to age acceleration to quantify the deviation of a cell from its expected predicted age. **b,** Predicted age as a function of actual age, with predicted ages computed for all cells by leaving out cells from one mouse and training aging clocks on the remaining cells to make predictions on the held-out cells and repeating this procedure for all mice. Heatmap colors represent density of predicted ages. Gray circles represent the median predicted age for an individual mouse. The line of best fit for the median predicted ages is shown in black. The Pearson correlation between predicted age and actual age for all cells is reported as *R* (with values of *R* > 0.7 in bold), and the Pearson correlation between median predicted age and actual age for all mice is reported as *r*. **c,** Dot plot comparing the predictive performance of single-cell aging clocks without spatial smoothing (“SingleCell”) with our single-cell spatial aging clocks (“SpatialSmooth”). Colors correspond to Pearson correlation between predicted age and actual age and the size of the dots are inversely related to the mean absolute error between predicted age and actual age. **d,** Density of predicted ages in an external 140-gene MERFISH dataset consisting of six coronal sections from three mice (3, 19, 25 months old) that shared 72 genes in common with the 300 genes used to train spatial aging clocks. Missing genes were imputed using SpaGE before spatial aging clock predictions were obtained. For statistical analysis, see Supplementary Table 12. See Supplementary Fig. 2a to compare with cross-validation results for the coronal sections dataset. **e,** Density of predicted ages across the young (<9 months) and old (>19 months) sagittal section samples. For statistical analysis, see Supplementary Table 12. See Extended Data Fig. 6c,d for predictions on regions not present in the coronal section training dataset (cerebellum, olfactory bulb). **f,** Summary of the training data used to build the spatial aging clocks and the generalization of spatial aging clocks to datasets containing different attributes. See Extended Data Fig. 5b,c, 6a-e, and Supplementary Table 12 for corresponding analyses.

The automatically selected features and number of genes used in each cell type-specific aging clock were different, ranging from 62 genes for T cells to 292 genes for astrocytes (Supplementary Table 10). Spatial aging clock genes with positive coefficients were enriched for many distinct biological processes (Extended Data Fig. 5e, Supplementary Table 11), while spatial aging clock genes with negative coefficients were generally enriched for differentiation and development processes (Extended Data Fig. 5f, Supplementary Table 11). In line with the observation that each cell-type-specific spatial aging clock selects different sets of genes, spatial aging clocks exhibited better performance within the same cell type than across different cell types (Extended Data Fig. 5g).

Spatial aging clocks generally produced accurate predicted ages in the same cell type across all subregions of the coronal brain sections (Extended Data Fig. 5h). Spatial aging clocks trained on individual subregions also exhibited good performance (albeit less good than that of the spatial aging clocks trained across all regions) (see Methods, Extended Data Fig. 5i), and they could likewise generalize to the same cell types in other subregions of the coronal brain sections (Extended Data Fig. 5j).

We evaluated whether these spatial aging clocks are generalizable to external datasets. Our spatial aging clocks robustly separated three ages (young, middle-aged, old) on an independent MERFISH dataset of coronal brain sections that we had previously generated using a panel of 140 genes^40^, 72 of which overlapped with the 300 genes selected in this study (see Methods) (Fig. 2d). The separation of the three ages in this external dataset was overall similar to what we observed for cross-validation within the coronal section dataset used to train the clocks (Supplementary Fig. 2a). Spatial aging clocks had a slightly lower performance for separating ages for some cell types in external datasets (and to some extent for cross-validation on the coronal section dataset) – likely due to a combination of low gene overlap with our dataset, relatively small magnitude of transcriptomic changes with age in certain cell types (e.g. neurons), and/or low cell numbers (e.g. neuroblasts and NSCs). Interestingly, the spatial aging clocks, which were trained on data from male mice, generalized to an external publicly available MERFISH dataset of coronal brain sections in female mice (juvenile, young, and old) with 75 genes that overlapped^22^ (Extended Data Fig. 6a) and to a corresponding single-nuclei RNAseq dataset of female mice (juvenile and old)^22^ (Extended Data Fig. 6b).

We found that our spatial aging clocks, which were trained on cells from coronal brain sections, also generalized to sagittal brain sections which include regions that are not present in coronal sections (e.g. olfactory bulb, cerebellum) (Fig. 2e, Extended Data Fig. 6c,d, Supplementary Fig. 2b), and to dissociated single-cell RNAseq dataset of the entire brain (containing additional brain regions) from male mice (young and old)^4^ (Extended Data Fig. 6e). Overall, spatial aging clocks could robustly separate young and old across most cell types in all external datasets evaluated (see Supplementary Table 12 for statistical analysis). Even in cases where the absolute predicted ages may be less accurate likely due to partial gene overlap with cell type-specific spatial aging clocks, the relative differences in predicted ages across age groups were generally consistent (see Supplementary Table 12 for statistical analysis). Thus, the large number of cells profiled per sample using MERFISH enables the development of specific and high-performing aging clocks, particularly for rare cell types such as ependymal cells, macrophages, T cells, and pericytes. These spatial aging clocks can generalize to independent cohorts of mice, to external spatial transcriptomics datasets with only partial gene overlap, across sex, across brain regions, and to other single-cell transcriptomics data modalities (Fig. 2f).

Overall, our spatial aging clocks represent a powerful tool for quantifying aging that could be used to systematically understand the influence of interventions on a tissue at a spatially resolved and cell type-specific level.

### Spatial aging clocks reveal distinct patterns of brain rejuvenation from exercise and partial reprogramming

Several interventions have been shown to restore aspects of brain function in old age^41–63^. Spatial aging clocks could provide an unbiased and high-throughput way to assess the impact of such ‘rejuvenating interventions’ across different cell types and regions of the brain. To test this possibility, we generated additional MERFISH spatial transcriptomics datasets in young and old mice in response to two rejuvenating interventions–voluntary exercise or ‘partial reprogramming’ (i.e. cyclic *in vivo* expression of the Yamanaka factors *Oct4*, *Sox2*, *Klf4*, *c-Myc* [OSKM]). We chose these two interventions because they have beneficial effects on the old brain, are thought to impact many diverse cell types, and likely have different modes of action (systemic for exercise vs. cell-intrinsic for partial reprogramming)^48,50,54,57,61,62^.

For exercise, we used 4 young (3 months) sedentary male mice, 4 old (19 months) sedentary male mice, and 4 old (19 months) male mice that voluntarily exercised using a running wheel for 5 weeks (based on published regimen for efficacy^54,64^) (Fig. 3a). For partial reprogramming, we used transgenic mice that allow whole-body cyclic expression of OSKM^62,65^ consisting of 4 young (5 months) male mice, 4 old (26-29 months) male mice, and 4 old (26-29 months) male mice that underwent cyclic induction of OSKM expression (see Methods for details) (Fig. 3a). We then used MERFISH to profile the panel of 300 genes with an identical setup as that used for the aging atlas (i.e. coronal sections) (Fig. 1a). In the exercise intervention dataset, we characterized more than 900,000 cells in total and they also clustered into 18 cell types across all conditions (Fig. 3b, Extended Data Fig. 7a). In the partial reprogramming intervention dataset, we characterized more than 1 million cells in total and they clustered into 15 cell types across all conditions (Fig. 3c, Extended Data Fig. 7b). T cells, B cells, and neutrophils were not identified in the partial reprogramming dataset, in line with dissociated single-cell datasets^62^ and likely due to lower abundance of infiltrating immune cells in this model. While aging led to significant changes in the proportion of several cell types in these datasets (e.g. oligodendrocytes and NSCs) (Extended Data Fig. 7c,d), as observed in the aging atlas (see Fig. 1c), exercise and partial reprogramming interventions did not have a significant effect on cell type proportion in old mice (Extended Data Fig. 7c,d), perhaps due to the relatively low effect and sample size.

**Figure 3:**
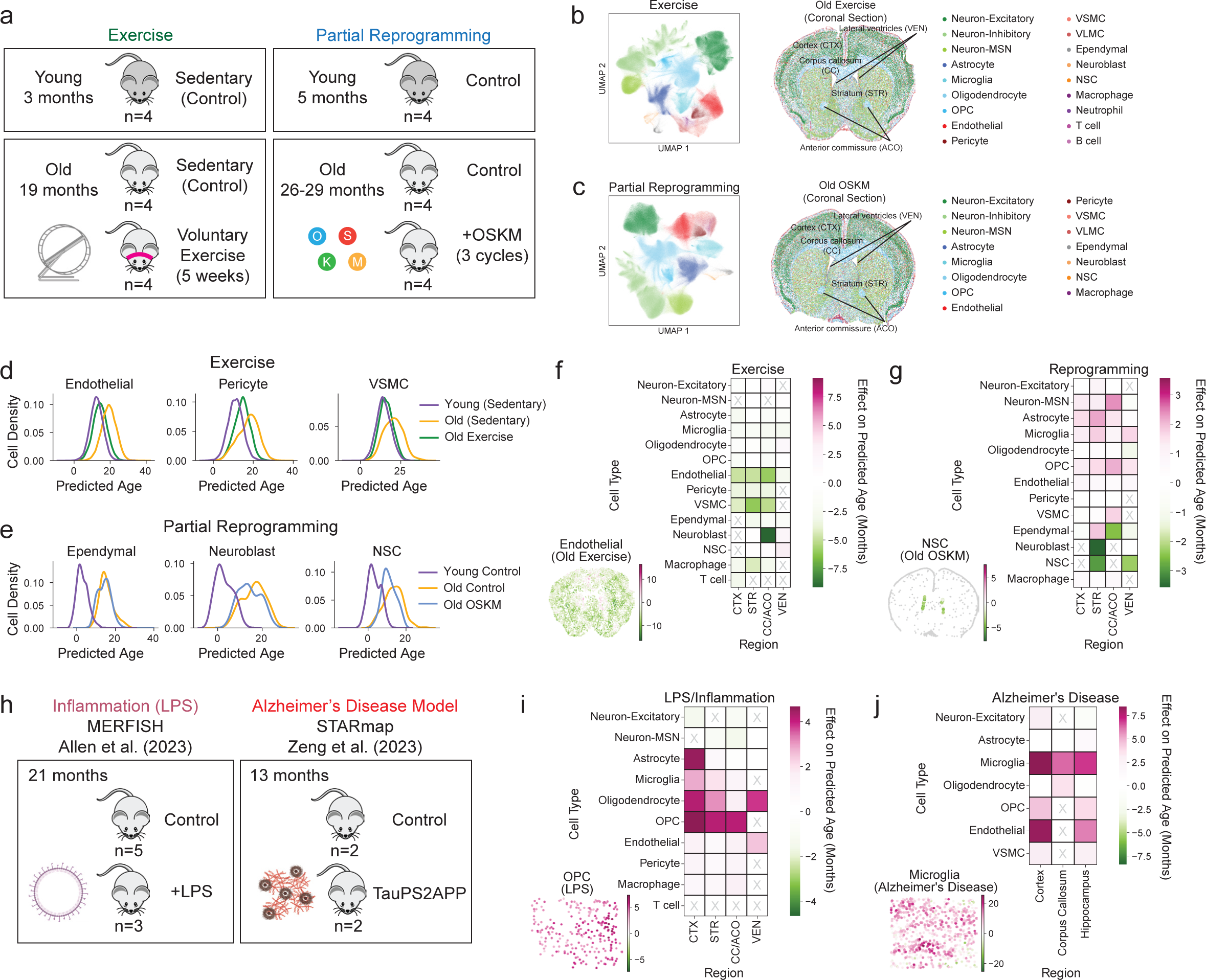
Impact of rejuvenation strategies, inflammation, and disease on spatial single-cell aging. **a,** Schematic illustration of the experimental cohorts for testing the effects of exercise and whole-body partial reprogramming as rejuvenating interventions. Four male mice were used per condition. **b,** UMAP visualization of all cells from the exercise study colored by cell type (left) and visualization under spatial coordinates for all cells in one representative coronal section from an old exercise mouse (right). Brain regions are annotated. **c,** UMAP visualization of all cells from the whole-body partial reprogramming study colored by cell type (left) and visualization under spatial coordinates for all cells in one representative coronal section from an old OSKM mouse (right). Brain regions are annotated. **d,e,** Density of predicted ages across different experimental conditions for spatial aging clocks corresponding to **(d)** three cell types rejuvenated by exercise and **(e)** three cell types rejuvenated by whole-body reprogramming. For statistical analysis, see Supplementary Table 12. **f,g,** Heatmaps showing the effect of rejuvenating interventions on predicted age for different cell types and regions, measured as the difference in median predicted age between intervention and control conditions for old mice, under **(f)** exercise and **(g)** whole-body partial reprogramming. Gray “X” denotes cell types and regions with insufficient numbers of cells (<50). Insets show spatial visualization of cells in an example intervention sample colored by their difference from the median predicted age in the control condition, with endothelial cells in **(f)** and NSCs (with small gray VLMCs) in **(g)**. Size of cells in each inset was manually chosen for visual purposes. **h,** Schematic illustration of cohorts from publicly available spatial transcriptomics datasets for systemic inflammation models (LPS injection) with MERFISH and Alzheimer’s disease (AD) models (TauPS2APP) with STARmap. **i,j,** Heatmaps showing the effect of adverse interventions on predicted age for different cell types and regions, measured as the difference in median predicted age between intervention and control conditions for **(i)** old female mice under systemic inflammatory challenge by LPS injection and **(j)** middle-aged male Alzheimer’s disease model mice. Gray “X” denotes cell types and regions with insufficient numbers of cells (<50). Insets show spatial visualization of cells in an example intervention sample colored by their difference from the median predicted age in the control condition, with OPCs in **(i)** and microglia in **(j)**. Size of cells in each inset was manually chosen for visual purposes. See also Extended Data Fig. 8a-e for additional profiling of adverse interventions with spatial aging clocks.

Importantly, we asked if our spatial aging clocks could uncover which brain cell types and regions experienced the greatest transcriptomic rejuvenation (i.e. exhibiting a younger predicted age with intervention than old control) by exercise and partial reprogramming.

Our spatial aging clocks indicated that the transcriptomes of several cell types were rejuvenated by exercise, including endothelial cells (median rejuvenation of 4.9 months), pericytes (median rejuvenation of 3.4 months), and vascular smooth muscle cells (median rejuvenation of 4.7 months) (Fig. 3d,f, see Supplementary Table 12 for statistical analysis). These strong rejuvenating effects on cells of the brain vasculature were present across multiple brain regions (except for the lateral ventricles) (Fig. 3f), consistent with a systemic effect of exercise. The striking impact of exercise on the brain vasculature may be linked to their exposure to circulating blood factors, which mediate some effects of exercise^42,55^, and to the generation of new blood vessels (vasculogenesis) in the brain in response to exercise^66^. Gene set enrichment analysis revealed that genes downregulated by exercise were significantly enriched for cell junction and focal adhesion processes only in endothelial cells in the rejuvenated regions but not in the lateral ventricles (see Supplementary Table 13), consistent with the region-specific endothelial cell rejuvenation by exercise. Neuroblasts also experienced region-specific rejuvenation by exercise (Fig. 3f). Indeed, while neuroblasts are present in the lateral ventricles and adjacent regions such as the corpus callosum, they only showed a strong rejuvenation by exercise near the corpus callosum (Fig. 3f), perhaps due to the presence of specific growth factors in this region. Thus, overall, exercise has a strong rejuvenating effect on the brain, consistent with previous studies^41,42,50,52,53,55^, and this beneficial effect is particularly pronounced in endothelial cells and neuroblasts in specific regions of the brain

Our spatial aging clocks also revealed that the transcriptomes of a few cell types were rejuvenated by partial reprogramming, including NSCs (median rejuvenation of 2.7 months) and neuroblasts (median rejuvenation of 2.8 months) (Fig. 3e,g, see Supplementary Table 12 for statistical analysis), consistent with the restoration of neural progenitors by partial reprogramming from dissociated single-cell RNAseq data^62^. Ependymal cells exhibited region-specific rejuvenation near the corpus callosum but not in the lateral ventricles or near the striatum/adjacent regions (Fig. 3g). In contrast, other cell types (medium spiny neurons, microglia, and glial cells) were prematurely aged across multiple brain regions in response to partial reprogramming (Fig. 3g). Hence, this regimen of partial reprogramming might have both beneficial and detrimental effects on the brain, depending on the cell type.

Overall, our spatial aging clocks reveal the specific cell types and regions most impacted by two different rejuvenating interventions, highlighting divergent–and potentially complementary–modes of rejuvenation.

### Spatial aging clocks reveal accelerated aging in models of inflammation, disease, and injury

In addition to studying the effects of rejuvenating interventions, we applied our spatial aging clocks to compare the effect of interventions known to be detrimental to the brain. We analyzed a publicly available MERFISH spatial transcriptomics dataset of female mice subjected to systemic inflammatory challenge following injection of lipopolysaccharide (LPS) as a model of accelerated aging^22^ (Fig. 3h). The dataset contained parts of the cortex (CTX), striatum (STR), and corpus callosum (CC)^22^. We also analyzed a publicly available STARmap spatial transcriptomic dataset (another approach with single-cell resolution) on brain sections containing the hippocampus from male middle-aged (13 months) Alzheimer’s mouse models (triple transgenic TauPS2APP)^30^ (Fig. 3h). In both datasets, we found that a subset of cell type annotations overlapped with the 14 cell types for which we built high-performance spatial aging clocks (10 cell types in the LPS dataset, 7 cell types in the Alzheimer’s mouse model dataset).

Our spatial aging clocks predicted that the transcriptomes of many cell types in the brain had accelerated transcriptomic aging (i.e. exhibited an older predicted age with intervention than age-matched control) in response to inflammation and disease.

Our spatial aging clocks revealed that several cell types, including glial cell types (astrocytes, oligodendrocytes, OPCs) and microglia exhibited accelerated aging in response to LPS (Fig. 3i, Extended Data Fig. 8a). The impact of LPS on the predicted age of astrocytes was highly specific to the cortical regions, while the impact of LPS on oligodendrocytes, OPCs, and microglia was present across multiple brain regions (Fig. 3i, see Supplementary Table 12 for statistical analysis). The global accelerated transcriptomic aging of oligodendrocytes and OPCs may be driven by their upregulated inflammation state under LPS condition (Extended Data Fig. 8b), in line with reported upregulation of inflammatory (activation) states of astrocytes and microglia in response to LPS^22^.

Our spatial aging clocks revealed that most cell types (microglia, neurons, and cells of the brain vasculature) exhibited accelerated transcriptomic aging in the Alzheimer’s disease model across several brain regions (Fig. 3j, Extended Data Fig. 8c, see Supplementary Table 12 for statistical analysis). The strong accelerated transcriptomic aging of microglia in Alzheimer’s disease is consistent with the substantial enrichment of disease-associated microglia in this particular model^30^ and the known microglial defects of Alzheimer’s disease^30,67–71^.

For both LPS and Alzheimer’s disease, application of our spatial aging clocks to samples from younger animals revealed similar patterns in accelerated transcriptomic aging of cell types, with the accelerated aging effects generally being more pronounced in older animals (Extended Data Fig. 8d,e). These observations suggest that older animals may be more susceptible to the accelerated aging effects of inflammation and disease.

Demyelination injury in the brain (for example in the context of multiple sclerosis) has recently been linked to accelerated brain aging in humans^72–74^. We applied our spatial aging clocks to publicly available datasets on two different multiple sclerosis mouse models—an *in situ* sequencing (ISS) dataset of global demyelination upon experimental autoimmune encephalomyelitis (EAE)^31^ and a MERFISH dataset of localized demyelination injury^29^. Our spatial aging clocks revealed strong accelerated transcriptomic aging across most cell types, particularly for microglia, in the global EAE model compared to control mice across all brain regions (Extended Data Fig. 8f). In contrast, in the case of localized demyelination injury, the greatest age acceleration was spatially restricted to the site of injury for multiple cell types (Extended Data Fig. 8g), suggesting that localized interventions can have a strong and specific effect in the region of intervention.

Hence, spatial aging clocks reveal cell type-specific and spatial deleterious effects in inflammation, neurodegenerative disease, and demyelination injury models, providing a high-resolution understanding that should serve as a foundation for developing targeted interventions, which has not been possible with aging clocks built on other data modalities. Overall, the spatial aging clocks can record rejuvenation and accelerated aging effects across very diverse interventions known to impact brain aging but acting via different mechanisms (e.g. exercise, partial reprogramming, disease and injury models), suggesting that these models can provide an integrative view of aging.

### Proximity effects of cell types on the aging of nearby cells

Could some cells influence the age of other cells around them? This question is particularly critical in the brain, given the complexity of the tissue and the potential for a cell to impact many different cell types around it. Spatial aging clocks enable systematic characterization of the effect of cell type proximity on single-cell aging *in situ*. We quantified the spatial proximity effect of a given “effector” cell type on the deviation from the expected predicted age (i.e. “age acceleration”, see Methods) of a “target” cell type. To this end, we computed the nearest distance to any effector cell for each target cell. “Near” target cells have a nearest distance to an effector cell that is less than the median neighbor distance within a given subregion (see Methods for cutoffs), and “Far” target cells have the largest nearest distances to an effector cell. To account for differences in cell proportions with age or across spatial regions (see Fig. 1c,e), we matched “Near” and “Far” target cells separately for each subregion, mouse, and age (Fig. 4a). We then aggregated these groups and compared the statistical difference in age acceleration of the “Near” and “Far” target cell groups to determine the proximity effect of the effector cell type on the target cell type. The variance in age acceleration was similar between “Near” and “Far” target cell groups (Extended Data Fig. 9a).

**Figure 4:**
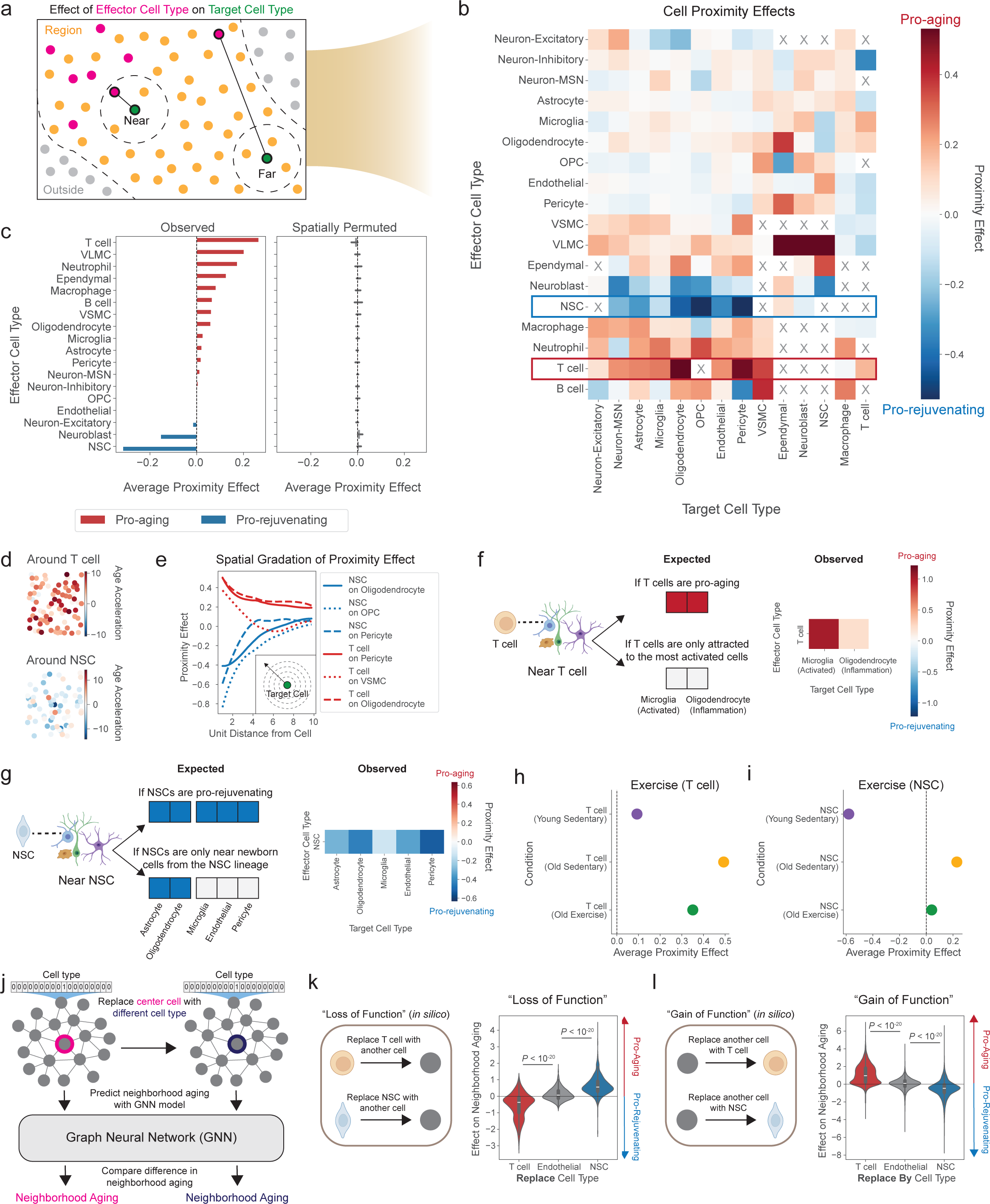
Proximity effects of cells on their neighbors during aging and rejuvenation. **a,** Schematic illustration of the matching process for determining “Near” and “Far” target cells to compute the proximity effect of effector cells on the age acceleration of target cells (coronal sections). Matching is controlled for region, age, and cell type. **b,** Heatmap showing the proximity effect, Cohen’s D of age acceleration of the “Near” target cells compared to the “Far” target cells, for different cell type proximity relationships. Rows correspond to the effector cell type and columns correspond to the target cell type, which experiences the proximity effect by the effector cells. Shown are the 14 target cell types with high-performing spatial aging clocks. “X” denotes proximity relationships for which there were insufficient cell pairings (<50) to compute a proximity effect (see Methods). Color bar is trimmed at top 2% absolute proximity effect values. **c,** Average proximity effect for a given effector cell type on all other target cell types ranked from most pro-aging (positive aging effect) to most pro-rejuvenating (negative aging effect) (left) and median average proximity effects of effector cell types computed after spatial permutation of all cells across each section (right). Error bars show 95% confidence interval across 20 spatial permutations. **d,** Spatial visualization of cells within 100 microns from an example T cell (top) and NSC (bottom) with colors corresponding to age acceleration. **e,** Spatial gradation of the proximity effect as a function of the unit distance from the target cell (see Methods) for the top two pro-aging proximity relationships with T cell as the effector cell type and the top two pro-rejuvenating proximity relationships with NSC as the effector cell type. Cutout shows scheme for spatial gradation analysis. **f,** Schematic showing expected heatmaps (left) for different T cell proximity effect scenarios and the observed heatmap (right) showing the T cell proximity effect on glial cell subtypes (activated/inflammation status, see Methods and Extended Data Fig. 4e). **g,** Schematic showing expected heatmaps (left) for different NSC proximity effect scenarios and the observed heatmap (right) showing NSC proximity effect on terminal cell types in the NSC lineage (astrocytes, oligodendrocytes) and cells not in the lineage (microglia, endothelial cells, pericytes) reproduced from (b). **h,i,** Average proximity effects of **(h)** T cells and **(i)** NSCs computed from all nearby cells in the exercise study for each of the three experimental conditions (Young Sedentary, Old Sedentary, Old Exercise). **j,** Graph neural network (GNN) models trained to predict neighborhood aging from local cell graphs with cell type features. These GNN models can be used to measure the effect of cell type perturbations on neighborhood aging and add a layer of causality. **k,l,** Effect of perturbing T cells, endothelial cells, and NSCs on neighborhood aging: **(k)** perturbations involving replacing the effector cell type with another cell type (“loss of function”) for >25,000 local cell graphs and **(l)** perturbations involving replacing another cell type with the effector cell type (“gain of function”) with >1,500 local cell graphs per effector cell type. The inner box corresponds to quartiles and the whiskers span up to 1.5 times the interquartile range of the effects across all cell networks. *P*-values computed using two-sided Mann-Whitney test to compare neighborhood aging perturbation effects for T cells or NSCs to that for endothelial cells.

After computing the proximity effect for all combinations of effector and target cell types and filtering for relationships that occur at sufficient frequency in the coronal section dataset (see Methods), we identified 214 distinct cell-cell proximity effects (Fig. 4b, Supplementary Table 14). Target cell types that were the most impacted by any effector cell type were cells in the lateral ventricles (ependymal cells, NSCs, neuroblasts), pericytes, and OPCs (Extended Data Fig. 9b). The proximity effects of effector cell types were generally similar in different brain regions, though microglia had a stronger pro-aging effect on multiple target cell types in the corpus callosum and anterior commissure (Extended Data Fig. 9c).

We were particularly interested in effector cells with the largest global impact on their neighbors. Notably, T cells had the strongest pro-aging average proximity effect (Fig. 4c,d), and this influence was especially clear on oligodendrocytes and pericytes (Fig. 4b). The pro-aging proximity effect of T cells (which are mostly cytotoxic in the aging brain, Supplementary Fig. 3) was most pronounced in older mice (Extended Data Fig. 9d). Surprisingly, NSCs (and neuroblasts) had the strongest pro-rejuvenating average proximity effect (Fig. 4c,d), and this effect was particularly evident on OPCs and pericytes (Fig. 4b). The pro-rejuvenating proximity effect of NSCs was most pronounced in younger mice (Extended Data Fig. 9d).

The effects of T cells and NSCs on nearby cells are robust. T cells remained the most pro-aging effector cell type and NSCs remained the most pro-rejuvenating effector cell type even when different definitions of the “Far” cell group were used (Extended Data Fig. 9e,f), when the spatial aging clocks and proximity effect analysis were restricted to independent cohorts of the coronal section dataset (Extended Data Fig. 9g,h), and when two non-spatial aging clocks were used (see Methods, Extended Data 9i,j). Importantly, we verified that, on average, T cells were the most pro-aging cell type and NSCs were the most pro-rejuvenating cell type in multiple external datasets (Extended Data Fig. 9k, Supplementary Table 15).

The observed proximity effects are unlikely to be confounded by cell type proportion changes with age (see Fig. 1c). To test this, we randomly permuted the spatial locations of all cells in each sample and then performed the same cell proximity effect analysis. In the spatially permuted data, the proximity effects of all effector cell types were close to zero, ruling out confounding variables like cell type proportion influencing the magnitude of the proximity effects (Fig. 4c). Moreover, these proximity effects are unlikely to be confounded by spillover of transcripts between nearby effector and target cells due to potential issues in segmentation. In a first approach to test this, we analyzed proximity effects based on predicted ages obtained from spatial aging clocks trained on a subset of genes after filtering out genes with higher estimated spillover rate (see Methods) (Extended Data Fig. 9l). In a second approach to test this, we performed area-restricted proximity effect analysis, which excludes cells within a small radius of the effector cells, for spatial aging clocks and two non-spatial aging clocks (see Methods, Extended Data Fig. 9m,n). These different approaches all corroborated that T cells have the most pro-aging proximity effect and NSCs have the most pro-rejuvenating proximity effect (Extended Data Fig. 9l,n).

As expected, the proximity effects of T cells and NSCs on neighboring cells decrease in magnitude over increasing distance between effector and target cells (Fig. 4e). Interestingly, however, the pro-aging effect of T cells on nearby cells generally persists over larger distances than the pro-rejuvenating effect exerted by NSCs (Fig. 4e, Extended Data Fig. 9o), perhaps indicating that longer range pro-aging T cell effects could be propagated by more diffusible factors or cascades of events (e.g. successive inflammation of oligodendrocytes).

Are T cells having a pro-aging effect on neighboring cells or are they attracted to already aged and inflamed cells (for example activated microglia)^75^? To distinguish between these two possibilities, we selected the most activated microglia and inflamed oligodendrocytes based on expression of activation and inflammation signatures respectively (see Methods). We observed similar activation/inflammation signature levels between the most activated microglia and inflamed oligodendrocytes that were near or far from T cells (Extended Data Fig. 9p), though activated microglia and inflamed oligodendrocytes were spatially closer to T cells at a higher frequency than other cell types (Extended Data Fig. 9q,r). Importantly, however, we observed that T cells still had strong pro-aging proximity effects on oligodendrocytes or microglia even after controlling for activation and inflammation status (Fig. 4f). Together, these results suggest that T cells promote the aging of nearby cells, independent of their activation/inflammation state, though T cells may also be attracted to activated microglia and inflamed oligodendrocytes in a feedforward loop.

Are NSCs having a pro-rejuvenating effect on neighboring cells or are cells near NSCs mostly originating from newborn cells (and thereby predicted to be younger)? To test this, we compared the NSC proximity effect across differentiated cell types in the NSC lineage (astrocytes and oligodendrocytes) and differentiated cell types not in the NSC lineage (microglia, endothelial cells, and pericytes) (Fig. 4g). NSCs exerted similarly strong pro-rejuvenating effects on all target cell types, including those from a different cell lineage altogether, suggesting that the NSC proximity effect is not limited to newborn cells in the NSC lineage (Fig. 4g).

Finally, we wondered whether these proximity effects can be modulated by interventions to rejuvenate the brain. Using our spatially resolved transcriptomics data collected from the exercise experiment (see Fig. 3a), we first confirmed that the spatial pro-aging proximity effect of T cells is higher in old sedentary mice than young sedentary mice and that the pro-rejuvenating effect of NSCs is less pronounced in old mice (Fig. 4h,i, Extended Data Fig. 9d). In the context of exercise, the pro-aging proximity effect of T cells is reduced in old mice when subjected to voluntary exercise (Fig. 4h). Similarly, the NSC proximity effect is shifted in a rejuvenating manner by voluntary exercise in old mice (Fig. 4i). Interestingly, this strong shift in the pro-rejuvenating NSC proximity effect with exercise occurred in the absence of intrinsic rejuvenation of NSCs by exercise (Fig. 3f), suggesting that proximity effects can provide a complementary perspective on intervention outcomes. By contrast, we did not observe substantial impacts on spatial proximity effects resulting from partial reprogramming (Extended Data Fig. 9s), perhaps because of the cell-intrinsic nature of this intervention. Together, these results indicate that interventions can modulate cell proximity effects. These observations further suggest that the rejuvenation of the aging brain by exercise may result at least in part from reducing the pro-aging proximity effect of T cells and boosting the pro-rejuvenating proximity effects of NSCs.

### *In silico* perturbations to test the effect of T cells and NSCs on their neighbors

A key step in understanding the causal role of specific cells on their neighbors is to study perturbations. To test the effect of *in silico* cell type perturbations involving T cells and NSCs on the aging of nearby cells, we used a deep learning approach. To this end, we trained a graph neural network (GNN) model on local cell graphs defined around center cells to predict neighborhood aging, defined as the average age acceleration of all nearby cells, using only the cell type and graph connectivity as features (see Methods, Fig. 4j). Using this GNN model, we performed both “loss of function” and “gain of function” *in silico* manipulations by permuting the cell type of the center cell and measuring the effect on neighborhood aging. We found that replacing a T cell by another cell (“loss of function” experiment) led to a general decrease in the neighborhood aging (Fig. 4k) while replacing an NSC by another cell resulted in a general increase in the neighborhood aging (Fig. 4k). Conversely, replacing any cell by a T cell (“gain of function” experiment) led to an increase in neighborhood aging (Fig. 4l) while replacing any cell by an NSC resulted in a decrease in neighborhood aging (Fig. 4l). As a specificity control, we verified that in both setups, the replacement or addition of endothelial cells (a neutral cell type) did not substantially affect the neighborhood aging (Fig. 4k,l). Together, these *in silico* manipulations indicate that T cells have a pro-aging effect on their neighbors while NSCs have a pro-rejuvenating effect on their neighbors.

### Potential mediators of T cell and NSC proximity effects

What are the potential mechanisms that mediate the pro-aging effect of T cells or the pro-rejuvenating effect of NSCs on nearby cells? To address this question, we leveraged computational tools and assessed global changes in gene expression and pathways in nearby cells. To augment the 300 genes measured through MERFISH, we performed uncertainty-aware spatial gene expression imputation, using a method we have recently developed, TISSUE^40^, as a wrapper around the SpaGE^76^ and Tangram^77^ imputation algorithms. This imputation approach consists of jointly mapping transcriptomes from our MERFISH study with single-cell RNAseq datasets collected from mice across multiple ages and containing the brain regions of interest (e.g. lateral ventricles, corpus callosum, striatum)^2,3^, followed by prediction of new gene expression profiles using the single-cell RNAseq dataset as reference and differential analysis of gene expression and gene signatures (see Methods) (Fig. 5a). Using cross-validation, we verified that the imputed gene expression values were positively correlated with the actual gene expression values and generally had small absolute prediction errors (Fig. 5b) (see Methods).

**Figure 5:**
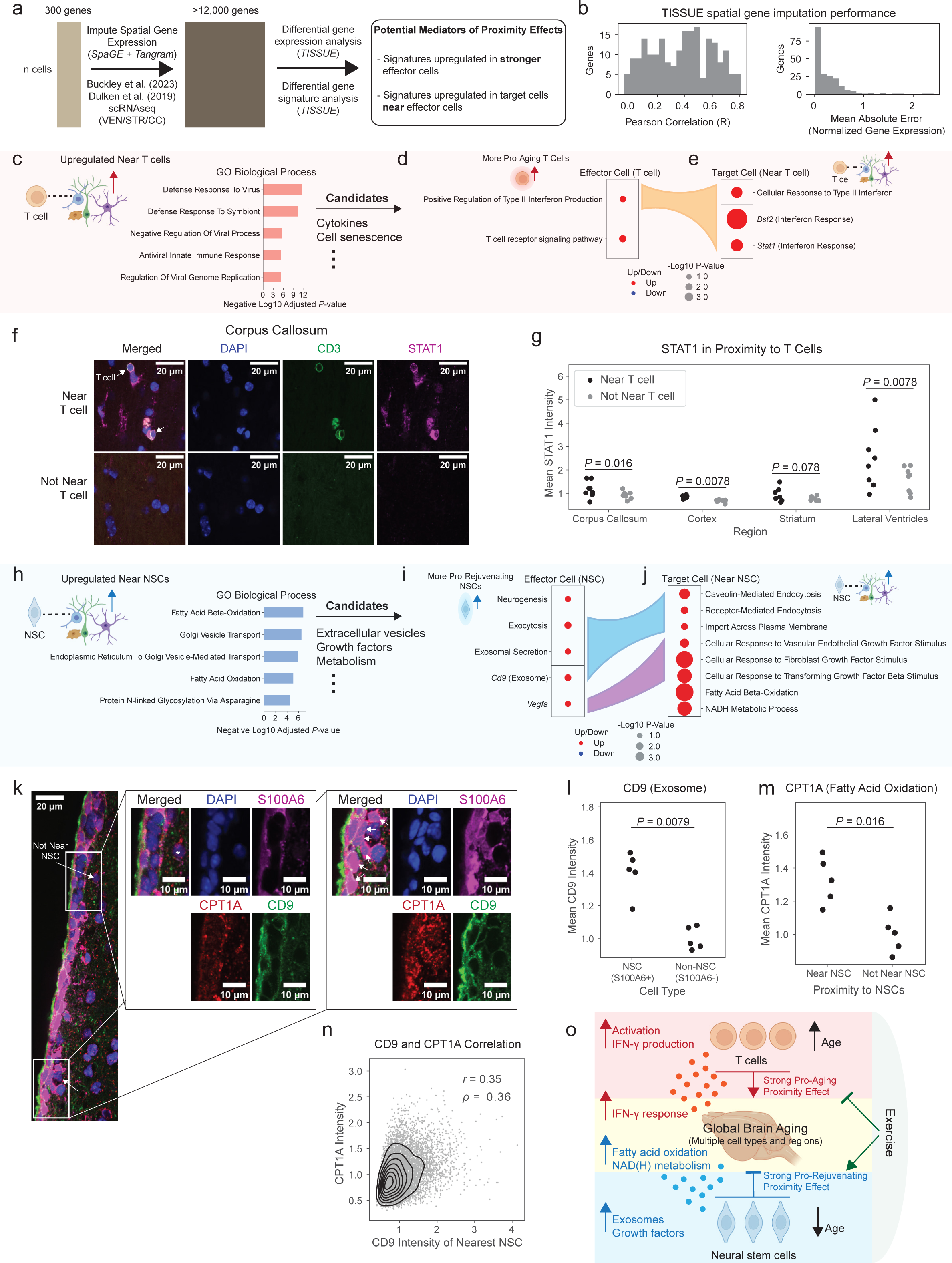
Molecular pathways underlying T cell and NSC proximity effects. **a,** Schematic illustration of the computational pipeline using SpaGE and Tangram algorithms for spatial gene expression imputation and using TISSUE for uncertainty-aware differential expression testing of imputed gene expression and gene signatures related to potential mediators of proximity effects. **b,** Histograms of gene-wise performance metrics for SpaGE imputation comparing predicted to actual gene expression using Pearson correlation (left) and mean absolute error (right). **c,** Bar plots showing top five most enriched GO Biological Processes for significantly upregulated genes in cells near T cells compared to cells far from T cells matched by region, age, and cell type. *P*-values computed from EnrichR pathway enrichment analysis**. d,e,** Dot plots showing the regulation of key GO Biological Process signatures (sum of imputed expression across all genes in set) and genes in the MERFISH panel for **(d)** imputed signatures in more pro-aging T cells compared to less pro-aging T cells; **(e)** imputed signatures and measured genes (*Bst2*, *Stat1*) in cells near T cells compared to cells far from T cells matched by region and cell type. Dot color corresponds to direction of regulation and dot size corresponds to negative log10 *P*-value (TISSUE two-sided t-test for GO Biological Process signatures and two-sided Mann-Whitney U-test for measured log-normalized gene expression comparisons). Colored flows represent sets of imputed signatures and genes that are related between different comparisons. **f,** Immunofluorescence staining image of a perfused mouse brain section from an old (28 months old) male mouse highlighting elevated STAT1 fluorescence (magenta) in cells that are near T cells (CD3^+^) (marked by arrows, top panels) compared to cells that are not near T cells (bottom panels) in the corpus callosum. Images were taken from the same brain section. **g,** Mean STAT1 intensity of cells that are near T cells compared to cells that are not near T cells across four independent brain regions (corpus callosum, cortex, striatum, lateral ventricles). Each dot represents the STAT1 intensities averaged across ∼25 cells that are near T cells or ∼190 cells that are not near T cells per region per section and then averaged across 3 sections for an individual mouse (see Methods). *P*-values computed with two-sided Wilcoxon signed-ranked test. **h,** Bar plots showing top five most enriched GO Biological Processes for significantly upregulated genes in cells near NSCs compared to cells far from NSCs matched by region, age, and cell type. *P*-values computed from EnrichR pathway enrichment analysis**. i,j,** Dot plots showing the regulation of key GO Biological Process signatures (sum of imputed expression across all genes in set) and genes in the MERFISH panel for **(i)** imputed signatures and measured genes (*Cd9*, *Vegfa*) in more pro-rejuvenating NSCs compared to less pro-rejuvenating NSCs; **(j)** imputed signatures in cells near NSCs compared to cells far from NSCs matched by region and cell type. Plot and statistical analysis parameters identical to (d,e). **k,** Immunofluorescence staining image of the perfused mouse lateral ventricle from a young (3.5 months old) male mouse with cutout highlighting example of elevated CPT1A fluorescence in cells that are near NSCs (S100A6^+^) with high CD9 fluorescence. In the full image, arrows within white rectangles label example cell near NSC and example cell not near NSC. Cutouts show merged and individual channels corresponding to image areas in the white rectangles. Arrows in the cutouts label NSCs. **l,** Mean CD9 intensity of NSCS (S100A6^+^) compared to non-NSCs (S100A6^-^). Each dot represents the CD9 intensities averaged across ∼290 NSCs or ∼380 non-NSCs per section and then averaged across 3 sections for an individual mouse (see Methods). *P*-value computed with two-sided Mann-Whitney U-test. **m,** Mean CPT1A intensity of cells that are near NSCs compared to cells that are not near NSCs. Each dot represents the CPT1A intensities averaged across ∼620 cells that are near NSCs or ∼120 cells that are not near NSCs per section and then averaged across 3 sections for an individual mouse (see Methods). *P*-value computed with two-sided Mann-Whitney U-test. **n,** Scatter plot of cells by CPT1A intensity as a function of the CD9 intensity of the nearest NSC (S100A6^+^) to that cell. All cells near NSCs from all individual animals (n=5) are shown. Gaussian kernel density estimate contours are shown in black. **o,** Summary of the main pro-aging and pro-rejuvenating proximity effects exerted by T cells and NSCs and their potential mediators.

To uncover potential mediators of T cell and NSC proximity effects, we used a two-step approach. First, we performed unbiased differential gene expression analysis using TISSUE^40^ on more than 12,000 imputed genes, with the goal of identifying common genes that were upregulated in cells near key effector cell types (T cells or NSCs) compared to cells far from the same effector cell type and matched by cell type and subregion (i.e. the same procedure as for calculating proximity effects). We verified that cells near T cells or NSCs did not include a cell type that was substantially more represented than all other cell types and that all 14 cell types with spatial aging clocks were present (Supplementary Table 16), and we checked that the variances in imputed gene expression were similar between the comparison groups (Extended Data Fig. 10a). In the second step, to better understand potential mechanisms of action, we performed targeted statistical comparisons of imputed gene signatures using TISSUE and gene expression in the MERFISH datasets for processes related to the most enriched terms from the unbiased analyses in both effector cell types (T cells or NSCs) and target cells (nearby cells). To this end, we divided effector cells into more effective (top 50%) vs. less effective (bottom 50%), i.e. more pro-aging T cells vs. less pro-aging T cells or more pro-rejuvenating NSCs vs. less pro-rejuvenating NSCs (see Methods). This approach allows us to augment the 300 gene panel of MERFISH and to screen for potential mediators of the T cell and NSC proximity effect.

### T cell proximity effect is associated with interferon signaling

Our differential gene expression analysis on over 12,000 genes revealed that cells near T cells experienced upregulation of genes associated with viral immune response (Fig. 5c), perhaps in response to interferon signaling. We therefore tested inflammation production and response pathways, notably interferon-γ which is involved in viral immune response as well as other candidates. T cells with greater pro-aging proximity effect exhibited increased expression of imputed gene signature for “positive regulation of type II interferon production” compared to T cells with less pro-aging proximity effect (*P* = 0.026, two-sided TISSUE t-test, Fig. 5d). Consistently, the more pro-aging T cells were more activated than their less pro-aging counterparts (Extended Data Fig. 10b), and T cell activation has been linked to increased interferon-γ production^78–80^. In comparison, T cells with greater pro-aging proximity effect did not show other candidate inflammatory production pathways, such as “positive regulation of type I interferon production” (*P* = 0.14, two-sided TISSUE t-test), “positive regulation of tumor necrosis factor production” (*P* = 0.47, two-sided TISSUE t-test), and “interleukin-6 production” (*P* = 0.22, two-sided TISSUE t-test). Consistent with the increased production of interferon-γ pathway in effector T cells, target cells near T cells exhibited concomitant increased expression of interferon-γ response genes (*Bst2*, *P* = 2.9×10^−14^; *Stat1*, *P* = 4.4×10^−5^; two-sided Mann-Whitney tests) in the MERFISH coronal section dataset (Fig. 5e). Target cells near T cells also showed increased expression of imputed gene signature for “cellular response to type II interferon” (*P* = 2.3×10^−4^, two-sided TISSUE t-test) compared to target cells far from T cells and matched by cell type and anatomic subregion (Fig. 5e).

Do conditions that lower interferon-γ in T cells modulate their proximity effects? We used the MERFISH dataset on LPS injection^22^, as LPS is known to dampen T cell activation and interferon-γ secretion^81–84^ (among other effects). In line with published findings, we observed significantly reduced scaled log-normalized expression of *Ifng* in the LPS condition compared to control (*P* < 10^−20^, two-sided Mann-Whitney U-test). Interestingly, lowering *Ifng* in T cells was associated with an attenuation of the T cell pro-aging proximity effect (Extended Data Fig. 10c). Together with the unbiased gene expression analysis, these data suggest a role for interferon-γ in mediating the T cell proximity effect.

We experimentally validated that the T cell proximity effect is associated with upregulated interferon response in nearby cells. To this end, we performed immunofluorescence staining on coronal brain sections of old mice (28 months) using antibodies to the T cell marker CD3 and the interferon-response marker STAT1, a marker that is also linked with inflammation-dependent aging^85^ (see Methods). We observed significantly higher STAT1 intensity in cells near T cells (CD3^+^ cells) compared to cells not near T cells, across the corpus callosum (*P* = 0.016, two-sided Mann-Whitney U-test), cortex (*P* = 0.0078, two-sided Mann-Whitney U-test), and lateral ventricles (*P* = 0.0078, two-sided Mann-Whitney U-test), and a trending increase in the striatum (*P* = 0.078, two-sided Mann-Whitney U-test) (Fig. 5f,g, Extended Data Fig. 10d). These findings experimentally validate the proximity effect of T cells on their neighbors and implicate interferon signaling as a key mediator of the T cell proximity effect.

### Pro-rejuvenating NSC proximity effect is associated with lipid metabolism

Our differential gene expression analysis on over 12,000 genes revealed that cells near NSCs experienced upregulation of genes associated with endocytic pathways and lipid metabolism (Fig. 5h). We therefore investigated components of the NSC secretome with known effects on lipid metabolism or endocytosis, which included extracellular vesicles/exosomes^86–88^ and growth factors^89–91^. NSCs with greater pro-rejuvenating proximity effect exhibited increased expression of imputed gene signatures for “exocytosis” (*P* = 0.014, two-sided TISSUE t-test) and “exosomal secretion” (*P* = 0.044, two-sided TISSUE t-test), increased expression of exosome marker *Cd9* (*P*=0.019, two-sided Mann-Whitney test), and trending increased expression of growth factor *Vegfa* (*P* = 0.059, two-sided Mann-Whitney test), which was the only NSC-secreted growth factor in the MERFISH panel (Fig. 5i). Extracellular vesicles/exosomes have been implicated in neurogenesis^92^ and could mediate the rejuvenating effect of NSCs on their neighbors. The increase in *Vegfa* is interesting in light of its effect on the maintenance of neurogenic niche cells^93–95^ and on longevity *in vivo*^96^. NSCs with greater pro-rejuvenating proximity effect also showed upregulation of pathways associated with cell proliferation and DNA replication (Extended Data Fig. 10e), which has been partly shown to be regulated by NSC-derived exosomes^97^. Concomitantly, cells near NSCs exhibited increased expression of imputed gene signatures for cell import and endocytosis (“caveolin-mediated endocytosis”, *P* = 1.9×10^−5^; “receptor-mediated endocytosis”, *P* = 7.2×10^−3^; “import across plasma membrane”, *P* = 0.017; two-sided TISSUE t-test) and cellular response to multiple growth factors (VEGF, *P* = 5.5×10^−4^; FGF, *P* = 3.3×10^−13^; TGFB1, *P* = 7.0×10^−10^; two-sided TISSUE t-tests) compared to cells far from NSCs, matched by cell type and anatomic subregion (Fig. 5j). Response to different growth factors can induce neurogenesis for repair and cognition^98–101^. Cells near NSCs also showed increased expression of a signature for NADH metabolism “NADH metabolic process” (*P* = 1.5×10^−9^, two-sided TISSUE t-test), which has been linked to fatty acid oxidation metabolism and more youthful brain states^102–104^ (Fig. 5j).

We experimentally validated, at the protein level, that the NSC proximity effect is associated with extracellular vesicle/exosomes in NSCs and upregulated fatty acid oxidation in nearby cells. To this end, we performed immunofluorescence staining on brain sections of young (3.5 months) mice using antibodies to the NSC marker S100A6^105^, the exosome marker CD9^106^, and the fatty acid oxidation marker CPT1A^107^, a marker that is linked to improved cell function^108^. We found that NSCs (S100A6^+^) had significantly higher CD9 intensity than other cell types (“non-NSCs”) in the lateral ventricles of young mice (*P* = 0.0079, two-sided Mann-Whitney U-test, Fig. 5k,l). Interestingly, cells that are near NSCs had significantly higher CPT1A intensity than cells that are not near NSCs in the lateral ventricles (*P* = 0.016, two-sided Mann-Whitney U-test, Fig. 5k,m). Importantly, there was a positive correlation between the CD9 intensity of NSCs and the CPT1A intensity in nearby cells (Pearson correlation *r* = 0.35, Spearman correlation ρ = 0.36, Fig. 5n). Together, these experimental validation results are consistent with the prediction that NSCs potentially mediate their pro-rejuvenating proximity effect through extracellular vesicles/exosomes that may affect nearby cells by upregulating fatty acid oxidation.

Together, our use of spatial aging clocks, deep learning models for *in silico* perturbations, gene imputation to screen for potential mediators, and experimental validation identifies cell proximity effects on aging and their potential modes of action (Fig. 5o). Our results suggest the following model (Fig. 5o): T cells have a strong pro-aging effect on nearby cells and infiltrate the brain during aging. These T cells express interferon-γ^2,29,109^, which may act on nearby cells in a pro-aging manner by inducing inflammatory responses. In contrast, NSCs have a strong pro-rejuvenating effect on nearby cells and decline in number with age. NSCs may secrete growth factors and extracellular vesicles/exosomes that act on nearby cells in a pro-rejuvenating manner, in part by modulating lipid metabolism.

## Discussion

Our study provides the highest-resolution spatiotemporal profiling of the aging mouse brain to date, enabling tracking of gene expression trends during aging in different regions and cell types. We use this dataset to generate the first spatial aging clocks (machine learning models that use spatially preprocessed gene expression to predict age) and, in turn, quantify the region-specific and cell type-specific effects of different rejuvenating interventions or disease models, providing a biological fingerprint for conditions that can potentially be used to identify synergistic combinations of rejuvenating interventions or targeted treatments for disease. Previous aging clocks have been built on different modalities (epigenetic, transcriptomics, proteomics)^110–122,60,123–128,3,129–139^, though in a spatially agnostic manner. While all clocks provide useful insight into the overall impact of interventions, only spatial aging clocks enable spatial and cell proximity discoveries.

The difference in predicted age (e.g. between young and old animals, between intervention conditions) produced by the spatial aging clocks are robust to experimental batch effects and partial overlap of gene panels; and they generalize across sex, different brain regions, and different single-cell transcriptomics technologies. How these spatial aging clocks apply to other tissues or species remains to be determined, but the flexible framework for building spatial aging clocks could be adapted to generate spatial aging clocks for other contexts. Spatial aging clocks should be generally helpful to rapidly assess the impact of experimental interventions on aging and other temporal processes (e.g. development, disease progression) at the spatial single-cell resolution. As recent studies reported spatial transcriptomic and epigenomic atlases of the young brain of mouse^18,34,140–144^, the tools we developed here could also apply to these types of datasets and other species, including humans. The development of new statistical methodologies to enhance the causality of aging clock features and improve the dimensionality of analyses as well as the development of experimental approaches allowing additional validation of spatial aging clock predictions could further broaden the usage of these models.

Notably, we use these spatial aging clocks to quantify cell proximity effects for the first time and show that they can be malleable to rejuvenating interventions such as exercise. These proximity effects are directional (i.e. effector cell acting on target cell) and confounding factors such as cell type proportion changes, age, animal, and anatomic region are controlled, which suggests that these proximity effects are primarily exerted by the effector cell. While determining a bona fide causal relationship underlying these proximity effects would be very challenging with current experimental techniques, we have pioneered a deep learning approach, based on graph neural networks, to measure effects on neighborhood aging resulting from *in silico* cell type perturbations. Similar approaches could be useful for identifying the effect of cell type or gene perturbations on other spatially localized readouts. A more systematic profiling of the whole brain across age and with a larger and unbiased gene panel could extend the characterization of cell proximity effects to more brain regions and cell types^33,34^, and may be particularly useful for developing an understanding of proximity effects in specific brain regions, such as different cortical layers, during aging. Deeper imaging for spatial transcriptomics may also allow the segmentation of long-range neuronal projections for greater resolution on the proximity effects of neurons. Although we identified potential mediating pathways for cell proximity effects using both spatial transcriptomics and immunofluorescence of protein markers, more targeted studies with functional cellular readouts could provide a better understanding of the mode of action and biological effects. Consideration of additional biological modalities may also offer a more systematic assessment of rejuvenation or accelerated aging. For example, different types of T cells have been shown to have either beneficial or detrimental effects on the brain^2,75,109,145–155^, particularly in response to injury or disease^29,156,157^. Disentangling the proximity effect of heterogeneous T cell populations in diverse regions could give a better understanding of neuroimmune interactions during aging. Finally, it will be interesting to determine how newborn cells in the brain (NSCs, neuroblasts) exert their pro-rejuvenating effects on their neighbors, whether extracellular vesicles are involved, and whether this could be implemented in other regions that do not contain stem cells. Ultimately, broader investigation into proximity effects and their mediators will be critical and may lead to new therapeutic strategies for improving brain resilience during aging.

## Acknowledgements

We thank members of the laboratory of A.B. and J.Z., especially Z. Huang, C. Bedbrook, J. Chen, and J. Na for technical discussion and assistance. We thank C. Bedbrook for independent checking of the code. We thank B. Engelhardt, L. Jerby, and M. Snyder for helpful discussion. We thank the Vizgen lab service team, especially J. He, Y. Cai, N. DiNapoli, and B. Wang for technical assistance. Elements of Figures 1-5 and Extended Data Figure 10 were created with BioRender.com. We would like to thank the Stanford Wu Tsai Neuroscience Microscopy Service. Support was provided by NSF Graduate Research Fellowship (to E.D.S.), P.D. Soros Fellowship for New Americans (to E.D.S.), Knight-Hennessy Scholars Program (to E.D.S.), NSF CAREER 1942926 (to J.Z.), grants from Silicon Valley Foundation (to J.Z.), Chan Zuckerberg Biohub–San Francisco Investigator program (to J.Z. and to A.B.), Chan Zuckerberg Initiative award (to A.B.), P01AG036695 (to A.B. and T.A.R.), R01AG071711 (to A.B.), the Milky Way Research Foundation (to A.B.), Simons Foundation grant (to A.B.), the Knight Initiative for brain resilience (to A.B.), and a generous gift from M. and T. Barakett (to A.B.).

## Author Contributions

E.D.S., J.Z., and A.B. planned the study. E.D.S. generated all datasets and performed all experiments and analyses, with contributions from other authors indicated below. M.H., O.Y.Z., and L.X. assisted with immunofluorescence experiments. O.Y.Z. assisted with sample collection for the exercise experiment. N.R. performed gene ontology term enrichment and regional gene expression change analysis and performed independent code checking. L.X. assisted with study design and sample collection for the partial reprogramming experiment. O.Y.Z., L.X., and P.N.N. assisted with sample collection for the aging dataset. L.L. and T.A.R. provided the exercise mouse cohort. E.D.S., J.Z., and A.B. wrote the manuscript, and all authors provided feedback.

## Methods

### Animals

All procedures involving mice were performed according to protocols approved by the Stanford University IACUC/AAPLAC (Protocol #8661) and VA Palo Alto Committee on Animal Research ACORP (LUO1736). For the aging cohort and exercise cohort, male C57BL/6JN mice were obtained from the National Institute on Aging (NIA) Aged Rodent colony. For the whole-body partial reprogramming cohort, male whole-body inducible OSKM (iOSKM) mice [*ROSA26(rtTA-M2); Col1a1(tetO-OSKM)*] (on a mixed background of the following strains: C57BL/6, B6D2F1, 129S4 and B6129SF1/J) were generated from the Jaenisch lab^158^ and obtained from the Jackson Laboratory (JAX 011004). Animals were housed in groups of 3-5 mice of the same age at the ChEM-H/Neuro vivarium (aging and partial reprogramming) or at the Veterinary Medical Unit at the Veterans Affairs Palo Alto Health Care System (exercise) for at least 3 weeks before any experiments or sample collection occurred.

### Exercise experiment

The exercise experiment included three groups of male C57BL/6JN mice: young sedentary animals (n=4, 3 months old), old sedentary animals (n=4, 19 months old), and old exercise animals (n=4, 19 months old). Sample sizes were selected to allow testing of statistically significant differences of animal-level attributes (e.g. cell type proportion) using the non-parametric two-sided Mann-Whitney U-test across groups. The animals in the old exercise group were provided with voluntary wheel running through individual housing for 5 weeks in polycarbonate cages with 12.7-cm-diameter running wheels (Lafayette Instrument, 80820), and monitored weekly for adequate running. Sedentary animals were individually housed in identical ages without running wheels.

### Whole-body partial reprogramming experiment

The whole-body partial reprogramming experiment included three groups of male iOSKM mice (on a mixed background of the following strains: C57BL/6, B6D2F1, 129S4 and B6129SF1/J): young control animals (n=4, 4.8-4.9 months old), old control animals (n=4, 25.6-29.2 months old), and old OSKM animals (n=4, 26.5-29.2 months old). Sample sizes were selected to permit testing of statistically significant differences of animal-level attributes (e.g. cell type proportion) using the non-parametric two-sided Mann-Whitney U-test across groups. Old animals were matched by age and body weight across control and OSKM. All animals (control and OSKM) were individually housed during the experiment. The animals in the old OSKM group were provided with three periods of cyclic induction of OSKM by doxycycline treatment, which consisted of doxycycline administration in the drinking water for two days (ON), followed by five days without doxycycline administration (OFF) and repeated for 3 weeks (ON-OFF-ON-OFF-ON) with animals sacrificed at the end of the last doxycycline administration treatment. Doxycycline (Fisher ICN19895505) was dissolved in drinking water (1 mg/ml) in amber water bottles to protect the solution from light and provided *ad libitum* to the animals in the old OSKM group.

### Sample collection

Animals were sacrificed to collect fresh frozen whole brain samples for MERFISH experiments. Animals were euthanized with 5 minutes of exposure in a CO_2_ chamber, and brains were removed and placed in a cryomold on ice and filled with pre-chilled Optimal Cutting Temperature (O.C.T.) Compound (Fisher Healthcare Tissue Plus, 4585) and then placed on dry ice. After O.C.T. solidified, the samples were moved to long-term storage at –80°C. RNaseZap (Invitrogen, AM9780) was used to disinfect all dissection tools before and after each animal. The first sample collection batch occurred on November 4, 2022 from 2:00-3:30pm PST and the second sample collection batch occurred on June 28, 2023 from 2:30-4:15pm PST. For the exercise experiment, mice were perfused with 15 mL of PBS with heparin sodium salt (50 U/ml) (Sigma-Aldrich, H3149-50KU) before sample collection. Sample collection for the exercise experiment occurred on June 6, 2022, and sample collection for the partial reprogramming experiment occurred on June 28, 2023.

### MERFISH 300 gene panel selection

We selected 300 genes to profile for the MERFISH experiments. Our selected 300 gene panel consists of 129 cell type and subtype markers (81 cell type markers, ranging from 1 to 8 markers per cell type; 48 subtype or function-related markers) as well as 181 genes of interest that are broadly related to brain aging or important biological pathways, with 10 genes shared across cell type and subtype markers and genes of interest (Supplementary Table 1).

The cell type and subtype marker genes included all markers in a suggested panel by Vizgen; markers determined from literature review focused on cells of the brain vasculature^4,159,160^, neural stem cells and neuroblasts^2,4,161,162^, and immune cells^2,4,163,164^. For these 129 cell type or subtype markers, we also included several neural stem cell and neuroblast markers obtained from integrated differential gene expression analysis across multiple single-cell RNAseq datasets of the adult mouse subventricular zone^2,3^. We included markers for the following cell types: excitatory neurons, inhibitory neurons, medium spiny neurons, astrocytes, microglia, oligodendrocytes, oligodendrocyte progenitor cells, endothelial cells, pericytes, vascular smooth muscle cells, vascular leptomeningeal cells, ependymal cells, neuroblasts, neural stem cells, macrophages, neutrophils, T cells, B cells, NK cells, mast cells, and dendritic cells.

For the aging-related genes, we included a set of 33 genes related to murine aging selected from the GenAge model organism database^37^ (Supplementary Table 1) to enrich for genes known to play a causal role in aging, which is helpful for building more causal aging clocks^38,39^. We further included genes related to T cell activity^2^, SVZ NSC heterogeneity^165^, endothelial heterogeneity^166^, meningeal lymphatics functions^160,167,168^, cellular senescence, immune response, stem cells, neurogenesis^169^, and vasculogenesis^170^. We also included sets of expert-curated genes pertaining to interesting cellular and organismal functions, some of which have emerging roles in the regulation of aging but are not well studied yet, including T cell signaling, reprogramming, cell adhesion/migration, lipid metabolism, and neuropeptide signaling. Finally, for a more unbiased set of genes, we included several differentially expressed genes between young and old mice for cell types across three single-cell transcriptomics datasets of the SVZ^2,3^ and two multi-region brain single-cell transcriptomics datasets^4,6^.

We ensured that all selected genes in the MERFISH panel were expressed in existing single-cell or single-nuclei RNAseq brain atlases^3,4,6^ and met technical constraints for minimizing optical clouding. This included limiting the total estimated gene expression using the Vizgen Gene Panel Design Portal to under 9,000 FPKM (total estimate at 7691 FPKM) and limiting the maximum estimated expression per gene using the Vizgen Gene Panel Design Portal to under 700 FPKM (maximum per-gene estimate at 452 FPKM). The complete gene panel, classification of markers, and rationale for inclusion are included in Supplementary Table 1.

### MERFISH imaging experiment

The MERFISH experiment was conducted through the Vizgen MERSCOPE technology lab service. Compared to other spatially resolved single-cell transcriptomics, Vizgen MERSCOPE technology has been shown to provide high specificity and sensitivity even with larger gene panel sizes^171^. Fresh frozen mouse brain samples were cut into 10-μm-thick sections on a cryostat at -20°C and placed onto a MERSCOPE slide (Vizgen 20400001). Both coronal and sagittal sections were taken at a depth to include the lateral ventricles, particularly the subventricular zone neurogenic niche, among other brain regions (cortex, striatum, corpus callosum). Two coronal sections were placed on each slide and paired to balance ages while sagittal sections were placed on their own slides. The tissue sections were fixed with 4% paraformaldehyde in 1x PBS for 15 minutes, washed three times with 5mL 1x PBS and incubated with 70% ethanol at 4°C overnight for tissue permeabilization. Samples were then stained for cell boundary using Vizgen’s Cell Boundary Kit (10400009), and later hybridized with a custom designed MERSCOPE Gene Panel Mix consisting of 300 genes (Vizgen 20300008) at 37°C incubator for 36-48 hours. Following incubation, the tissues were washed with 5mL Formamide Wash Buffer at 47°C for 30 minutes, twice and embedded into a hydrogel using the Gel Embedding Premix (Vizgen 20300004), ammonium persulfate (Sigma, 09913-100G) and TEMED (N,N,N’,N’-tetramethylethylenediamine) (Sigma, T7024-25ML) from the MERSCOPE Sample Prep Kit (10400012). After the gel mix solution solidified, the samples were cleared with Clearing Solution consisting of 50μL of Proteinase K (NEB, P8107S) and 5mL of Clearing Premix (Vizgen 20300003) at 37°C overnight. After removing clearing solution, the sample was stained with DAPI and Poly T Reagent (Vizgen 20300021) for 15 minutes at room temperature, washed for 10 minutes with 5mL of Formamide Wash Buffer, and then imaged on the MERSCOPE system (Vizgen 10000001). A fully detailed, step-by-step instruction on the MERFISH sample prep the full protocol is available at https://vizgen.com/resources/fresh-and-fixed-frozen-tissue-sample-preparation/. Full Instrumentation protocol is available at https://vizgen.com/resources/merscope-instrument/.

### Cell segmentation and MERFISH data preprocessing

Segmentation of cells was performed using Cellpose (1.0.2) through Vizgen’s lab service. Cell segmentation was implemented on images using nuclear staining (DAPI) and cytosolic staining (Poly T). Transcripts were allocated using these cell segmentations by summing across seven z-stacks, accounting for both nuclear and cytosolic (soma) transcripts. Quality control statistics were computed using the Vizgen post-processing tool for cell segmentation (Supplementary Table 2).

For preprocessing, we performed initial cell filtering separately for each MERFISH experiment (consisting of either two coronal sections on the same slide or one sagittal section). For each experiment, we removed putative doublets using Scrublet^172^ and a doublet score cutoff of 0.18. We then filtered out all cells with segmentation volume less than or equal to 100 or greater than or equal to three times the median cell volume. We also filtered out all cells with less than or equal to 20 counts and/or less than or equal to 5 genes with non-zero expression. To correct for potential different segmentation sizes, we divided the raw transcript counts obtained for each cell in the MERFISH dataset by the volume of the corresponding segmentation. After combining all experiments into an integrated dataset, we then filtered out all cells in the top 2% highest and top 2% lowest total expression. Statistics associated for the aforementioned cell filtering steps and for additional steps after clustering can be found in Supplementary Table 3. To obtain log-normalized gene expression values, we normalized the total gene expression for each cell to 250 and log-transformed the expression with an added pseudocount. This procedure was performed separately for the aging (coronal), aging (sagittal), exercise, and partial reprogramming cohorts.

### Cell type clustering and identification

For clustering, we converted each log-normalized gene expression value to a z-score using scanpy.pp.scale with max_value=10 in the scanpy package^173^. We performed Leiden clustering using scanpy.tl.leiden with resolution=0.5 for the initial clustering and default settings otherwise. We obtained batch-balanced nearest neighbors graph using BBKNN (scanpy.external.pp.bbknn) and then used this neighbors graph to generate a UMAP visualization of all cells (scanpy.tl.umap). For the partial reprogramming experiment, which only involved one batch of MERFISH data, we used scanpy.pp.neighbors with n_pcs=20 and n_neighbors=15 instead of BBKNN. To annotate cell types, we manually labeled each cluster based on cell type expression patterns as observed across two orthogonal data visualization modalities (the UMAP visualization and a heatmap of cell type markers) to reduce errors in cell type annotation resulting from dimensionality reduction distortions^174,175^. For clusters that expressed markers from multiple cell types, we performed successive Leiden clustering on those clusters until unique cell types could be annotated (see Supplementary Table 5). This procedure was performed separately for the aging (coronal), aging (sagittal), exercise, and partial reprogramming cohorts. A detailed description of the cell type markers and Leiden clustering resolutions for each dataset and cell type can be found in Supplementary Table 5. Although a small set of markers for some rare immune cell types (NK cells, mast cells, dendritic cells) were included in our MERFISH panel (Supplementary Table 1), we were unable to identify these cell types in any of the datasets (Supplementary Table 5), likely due to their low abundance. NK cells, mast cells, and dendritic cells were also not identified in several existing spatial transcriptomics studies of adult mouse brain^22,29–32^. However, NK cells and dendritic cells were identified in a large-scale and high-resolution spatial transcriptomic profiling of the whole adult mouse brain^33^, although at 3-4 times lower abundance than the rarest immune cell types identified in our datasets (T cells and B cells). In addition, in the partial reprogramming dataset, we were unable to identify other rare immune cell types (T cells, B cells, neutrophils), consistent with our dissociated single-cell RNAseq datasets^62^ and likely due to the lower abundance of these cell types in the partial reprogramming mouse model.

### Spatial region and subregion clustering and annotation

To identify anatomical regions across the MERFISH datasets and assign region and subregion labels to each cell, we adapted a semi-supervised approach for clustering and annotating region labels from cell type composition of local neighborhoods around each cell^22^. For a given cell, we computed the cell type abundances for each cell within 100um distance from the given cell. Then, we performed principal component analysis on the matrix consisting of the cell type abundance profiles for each cell, applied k-means clustering (k=25), manually visualized and merged clusters to obtain seven subregion annotations (CC/ACO, CTX_L1/MEN, CTX_L2/3, CTX_L4/5/6, STR_CP/ACB, STR_LS/NDB, VEN), and finally merged subregion annotations to obtain four region annotations (CC/ACO, CTX, STR, VEN). While we observed some variability in the subregion annotations between samples, there was general consistency in the four region annotations. We verified the expression of cortical layer markers in the three subregions of the cortex (Extended Data Fig. 3c,d), but we were unable to annotate each of the six known cortical layer individually through the clustering procedure, perhaps due to the low number of cortical layer markers. This region and subregion clustering and annotation procedure was performed separately for the aging (coronal), exercise, and partial reprogramming cohorts.

### Cell type composition analysis

We computed cell type proportions for a given sample by dividing the number of cells of each cell type by the total number of cells in the sample. For regional cell type proportions, we divided the number of cells of each cell type in that region by the total number of cells in that region. Pearson correlation, 95% confidence interval for the correlation, and *P*-value for association between cell type proportion and sample age was computed using scipy.stats.pearsonr. Linear regression of cell type proportion on sample age with 95% confidence interval was computed using seaborn.regplot. To compute statistical significance of differences in cell type proportions across categorical conditions, we used the two-sided Mann-Whitney U-test.

### Increasing and decreasing gene expression with age analysis

For a given cell type, to identify genes that changed in expression with age, we computed the Spearman correlation between age and pseudobulk gene expression across samples in the coronal section dataset. The pseudobulk gene expression was computed as the mean log-normalized gene expression across all cells of the same cell type within a sample. For each gene, we obtained the Spearman correlation, the associated *P*-value, and the lower and upper bounds of a 95% confidence interval for the correlation. We classified genes as “Increasing” if they had Spearman correlation greater than 0.3 and with the lower bound of the 95% confidence interval greater than 0.0. We classified genes as “Decreasing, if they had Spearman correlation less than -0.3 and with the upper bound of the 95% confidence interval less than 0.0. To reduce false positives resulting from transcript spillover due to segmentation, we constrained our analysis to the 220 genes with less than 5% estimated spillover based on an internal metric using gene expression variations influenced by local cellular composition.

### Gene ontology enrichment analysis

We performed gene ontology (GO) enrichment analysis to determine biological processes that were enriched in different sets of genes. For genes that increase or decrease in expression with age that were identified in “Increasing and decreasing gene expression with age analysis”, we performed GO biological process enrichment analysis separately for each set of genes and for each cell type. For genes in different spatiotemporal gene expression trajectory clusters for oligodendrocytes in the corpus callosum/anterior commissure (CC/ACO) region (see “Spatiotemporal gene expression trajectory analysis” below), we performed GO biological process enrichment analysis for all genes present in each of the nine trajectory clusters separately after filtering out genes with greater than 5% estimated spillover based on an internal metric using gene expression variations influenced by local cellular composition. For GO enrichment analysis on genes used by the spatial aging clocks, we selected positive coefficient clock genes (up to 50 genes with the largest positive coefficients) and negative coefficient clocks genes (up to 50 genes with the largest negative coefficients) and performed GO biological process enrichment analysis separately for each set of genes. For GO enrichment analysis on differentially expressed genes in endothelial cells in response to exercise in old mice, we selected genes that significantly increased with exercise (increased in old exercise compared to old sedentary with *P* < 0.05 from two-sided Mann-Whitney U-test) and genes that significantly decreased with exercise (decreased in old exercise compared to old sedentary with *P* < 0.05 from two-sided Mann-Whitney U-test) and performed GO biological process enrichment analysis separately for each set of genes.

We performed GO biological process enrichment analysis by selecting genes for each analysis (as described above), and using all other genes measured by MERFISH as background. GO enrichment was performed with topGO^176^ (R package version 2.54.0) using Fisher’s exact test for all Biological Process terms.

### Regional gene expression changes with age

To compare different anatomic regions and subregions by the magnitude of gene expression changes with age, we selected the five youngest and five oldest mice in the data. For each cell type, we determined the minimum number of cells present across each of the mice and regions, excluding the lateral ventricles due to their low cell number. We then downsampled cells for each mouse and region, without replacement, to that minimum number, such that after sampling, all combinations of mouse and region had the same number of cells. We then excluded cell types in which the number of cells per mouse and region was less than 20. We calculated the transcriptional profile for each mouse and region by averaging across the volume-normalized expression of all cells from that mouse in that region, normalizing this profile to sum to 250, and performing a log transformation with an added pseudocount. To determine the change between old and young mice in a specific region, we subtracted the mean profile of the 5 young mice from the mean profile of the 5 old mice and averaged the absolute value of this difference across all genes after filtering out genes with greater than 5% estimated spillover based on an internal metric using gene expression variations influenced by local cellular composition. We repeated this process 20 times, each time sampling different cells.

### Spatiotemporal gene expression trajectory analysis

For each cell, we divided the raw transcript counts by the segmentation volume of the cell and then normalized by total count determined using the default settings of scanpy.pp.normalize_total. For each combination of cell type, anatomic subregion, and gene; we computed a vector of length 20 (trajectory) containing the mean expression of that gene in each age conditioned on a specified subregion and cell type. If the given cell type and subregion combination was not present in at least 70% of profiled ages, we classified that trajectory as “missing”. We performed last-value-carried-forward imputation to fill in missing values within trajectories that passed this threshold. Each trajectory was standardized by centering and scaling to unit variance with sklearn.preprocessing.StandardScaler^177^. Then, we performed k-means clustering using sklearn.cluster.KMeans with n_clusters=9, random_state=444, n_init=’auto’, and the matrix with each row corresponding to a scaled gene expression trajectory as input. The parameters of the clustering were selected to maximize the number of clusters while also maintaining qualitatively distinct trends in each cluster (Extended Data Fig. 4d,e). For each cluster of gene expression trajectories, we visualized the smoothed median and interquartile range of the scaled expression values. Smoothing was done with interpolating B-splines using scipy.interpolate.BSpline with s=20. We manually annotated each cluster based on the qualitative expression patterns. To visualize representative gene expression trajectories for each cell type, we performed the same smoothing procedure separately for each cell type-specific subset of trajectories from the clusters. We developed this trajectory clustering approach instead of using parametric methods (e.g. polynomial fitting) to provide a more unbiased characterization of different trajectories.

### SpatialSmooth soft pseudobulking procedure

For each cell type, we build spatial graphs connecting each cell with its 20 nearest neighbors by Euclidean distance and of the same cell type^178^. Spatial aging clock performance was generally robust to the choice of the number of nearest neighbors (*k*) in building this graph (Extended Data Fig. 5a). We computed a L1-normalized adjacency matrix representing the spatial graph. Then, the SpatialSmooth algorithm propagates gene expression features across the cell type-specific spatial graph by iterating the update equation until convergence (i.e. *X*_*t*+1_ ≈ *X*_*t*_):

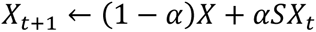

Where *X* is the initial gene expression matrix (cells as rows, genes as columns), *⍺* is the normalized adjacency matrix, and *⍺* is the smoothing parameter. We set *⍺* equal to 0.8. We set the convergence to be 30 iterations at maximum with a tolerance of 0.01, for which convergence is reached if ‖*X*_*t*+1_− *X*_*t*_‖_∞_ is less than the tolerance or 30 iterations has elapsed. For rare cell types (e.g. T cells, B cells, neutrophils), SpatialSmooth generally converged before 30 iterations. For common cell types (e.g. oligodendrocytes, microglia, astrocytes), SpatialSmooth was generally performed for 30 iterations. After convergence at step *t* = *T*, we use the smoothed spatial gene expression matrix *X*_*T*_ as input for training the aging clocks.

### Training and cross-validated evaluation of spatial aging clocks

For training spatial aging clocks, we performed the SpatialSmooth procedure on the log-normalized gene expression to obtain smoothed spatial gene expression matrices for each cell type independently. Then, for each cell type, we fitted a pipeline consisting of standardization of gene features followed by lasso regression model to predict sample age from a cell’s gene expression profile and used sklearn.linear_model.LassoCV to select optimal hyperparameters with cv=5, n_alphas=20, max_iter=10000. We refer to this entire pipeline from SpatialSmooth to age prediction as the ‘spatial aging clock’. To avoid explicit conditioning of age prediction on spatial information, the spatial aging clocks only leverage the spatial context to process the input data via SpatialSmooth.

For cross-validated evaluation of spatial aging clocks, we held out a single sample/age as the test set and kept the remaining samples/ages as the train set. SpatialSmooth was performed separately for the train and test sets. We fitted the lasso regression pipeline to predict age on the train set and used the model to obtain predicted ages on the test set. We repeated this procedure across all samples/ages to obtain predicted ages for all cells in the study. Performance was evaluated by Pearson’s correlation (*R*) and mean absolute error between the age and predicted ages of individual cells obtained from cross-validation. We also computed the Pearson’s correlation (*r*) between the age and median predicted ages of individual mice obtained from cross-validation. For the subregion-specific aging clocks, we trained and evaluated the models using the same settings except with 5 nearest neighbors for SpatialSmooth and restricted to only cells in each subregion for training.

### Visualization of spatial aging clock predictions

We used two approaches for visualizing the predicted ages obtained from the application of spatial aging clocks, either through cross-validation on the coronal section dataset or directly through validation on an external dataset.

For datasets with relatively uniform distribution of many ages across lifespan (i.e. the coronal section dataset), we used a correlation plot visualization consisting of a two-dimensional histogram of cell frequencies across bins defined by predicted age and actual age that is visualized as a heatmap, a scatter plot of the median predicted age of cells across each sample as a function of the actual age, and a line of best fit for the median predicted ages as a function of actual age is shown in black and computed using numpy.polyfit with deg=1. This type of visualization emphasizes the quality of median predicted age at the sample level across many different actual age values. Generally, in this visualization, the range of predicted ages will be larger than the range of actual ages due to heterogeneity in the predicted ages but not in the actual ages of cells from the same mouse. This may be especially pronounced for highly abundant cell types like excitatory neurons. Predicted ages obtained from cross-validation using a leave-mouse-out approach will exhibit some regression to the mean age (i.e. cells from older mice predicted to be younger and cells from younger mice to be predicted to be older) due to different mean actual ages in each of the cross-validation training datasets.

For datasets with bimodal distribution of ages (i.e. the sagittal section dataset) or with three or fewer distinct age groups (i.e. all external datasets), we used a density plot visualization of predicted ages of cells across different age groups or different experimental conditions. A kernel density estimate was constructed for each group of predicted ages using seaborn.kdeplot with default settings. This type of visualization emphasizes the distribution of predicted ages in a small number of groups at the cell level. For the coronal sections dataset and sagittal sections dataset, we used both types of representations.

### Application of spatial aging clocks on external datasets

To apply the spatial aging clocks to predict age for cell gene expression profiles in external spatial transcriptomics datasets, we applied the following general procedure. First, we filtered the external dataset to only include genes present in our MERFISH panel of 300 genes and only cell types also represented among the spatial aging clocks. Then, we normalized and log-transformed the raw expression values using the same approach as for our MERFISH data and applied SpatialSmooth (*⍺*=0.8) to the log-normalized values for each cell type independently. For clock genes that are not present in the external dataset, we use the training data for the clock as reference data in the SpaGE algorithm (n_pv=15)^76^ to impute the expression of the missing genes. Negative imputed values were clipped to zero. We performed imputation for genes from our MERFISH panel that were missing from these datasets (228 genes in the 140-gene MERFISH coronal section dataset, 5 genes in the single-nuclei RNAseq dataset, 36 genes in the single-cell RNAseq dataset, 225 genes in the LPS dataset, 128 genes in the Alzheimer’s mouse model dataset, 240 genes in the global demyelination through experimental autoimmune encephalomyelitis (EAE) dataset, 236 genes in the localized demyelination injury dataset). Finally, for each cell type, we applied the corresponding spatial aging clock (fitted standardization and lasso linear model) to generate predicted ages from the smoothed spatial gene expression values. For single-nuclei RNAseq datasets, which lack spatial information, we used the pseudobulk approach from a previous model^3^ with 20 cells contributing to each pseudocell instead of SpatialSmooth. For all evaluations, we also quantified the magnitude of difference in median predicted age (in units of months) and the statistical significance of this difference by performing two-sided Mann-Whitney U-tests on both the predicted ages of all cells between different pairs of age groups and the predicted ages of all mice (computed as the median predicted age) between different pairs of age groups. These statistics are reported in Supplementary Table 12. In some cases, imputation resulted in biased age predictions, but the differences in predicted age across ages and conditions were generally robust. In applying spatial aging clocks with imputation, we recommend comparing the predicted age to known ages in the dataset to calibrate interpretations. Generally, we observed lower spatial aging clock performance for cell types with low transcriptomic changes with age such as neurons (see Fig. 1f) or those with limited marker genes in external datasets such as neuroblasts (see Fig. 2d). Results were also generally consistent for all applications of the spatial aging clocks without SpaGE imputation and when using spatial aging clocks trained the 220 genes with less than 5% estimated spillover based on an internal metric using gene expression variations influenced by local cellular composition (Supplementary Table 12).

### Age acceleration calculation

We computed age acceleration for the predicted age of each cell to measure the deviation from its expected predicted age (i.e. the average of predicted age across all cells from a given cell type and animal). For each sample *k* ∈ {1, …, *K*} and cell type *p* ∈ {1, …, *P*}, we define the set of cells belonging to both sample *k* and cell type *p* as *Q*, and the age acceleration for each cell *i*∈ *Q* is defined as:

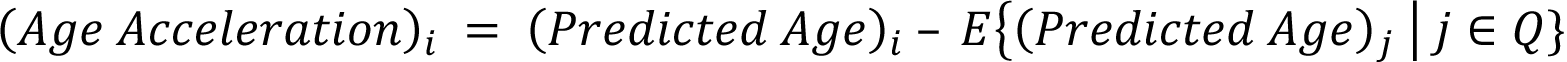

### Single-nuclei RNAseq aging data processing

We applied our spatial aging clocks to publicly available single-nuclei RNAseq data on the cortex and striatum of juvenile (0.93 months) and old (20.93 months) female C57BL/6J mice^22^. We downloaded processed data objects containing the scaled log-normalized gene expression from https://cellxgene.cziscience.com/collections/31937775-0602-4e52-a799-b6acdd2bac2e and mapped several cell types to our cell type classifications. We modified the preprocessing and imputation steps outlined in “<u>Application of spatial aging clocks on external datasets”</u> to account for the scaled log-normalized expression being used as input. Predicted ages were highly consistent when using a scaled log-normalized coronal section dataset for imputation.

### Single-cell RNAseq aging data processing

We applied our spatial aging clocks to publicly available single-cell RNAseq data on whole brain tissue (without hindbrain regions) of young (2-3 months) and old (21-22 months) male C57BL/6J mice^4^. We downloaded processed datasets from https://portals.broadinstitute.org/single_cell/study/aging-mouse-brain and mapped several cell types to our cell type classifications.

### 140-gene MERFISH aging data processing

We applied our spatial aging clocks to our 140-gene MERFISH spatial transcriptomics data on whole brain coronal sections of young (0.93 months), middle-aged (5.58 months), and old (20.93 months) male C57BL/6JN mice^40^. We mapped several cell types to our cell type classifications.

### MERFISH aging and LPS data processing

We applied our spatial aging clocks to publicly available MERFISH spatial transcriptomics data on the cortex and striatum of female C57BL/6J mice in juvenile (0.93 months), young (5.58 months), old (20.93 months), and LPS-injected conditions^22^. We downloaded processed data objects containing the scaled log-normalized gene expression from https://cellxgene.cziscience.com/collections/31937775-0602-4e52-a799-b6acdd2bac2e and mapped several cell types to our cell type classifications and used existing region annotations. We modified the preprocessing steps outlined in “<u>Application of spatial aging clocks on external datasets”</u> to account for the scaled log-normalized expression being used as input. Predicted ages were highly consistent when using a scaled log-normalized coronal section dataset for imputation. We included all control (aging) and LPS-injected animals in our analyses.

### STARmap Alzheimer’s disease data processing

We applied our spatial aging clocks to publicly available STARmap PLUS spatial transcriptomics data on the cortex and hippocampus of male TauPS2APP (AD model) and non-transgenic control mice across two ages (8 and 13 months)^30^. We downloaded processed data from https://singlecell.broadinstitute.org/single_cell/study/SCP1375 and mapped several cell types to our cell type classifications and used existing region annotations. We included all control and AD animals in our analyses. The percentage of zero counts was higher in this dataset (94%) than in the coronal section dataset used for training the spatial aging clocks (70%).

### *In situ* sequencing global demyelination data processing

We applied our spatial aging clocks to publicly available *in situ* sequencing (ISS) spatial transcriptomics data on whole brain coronal sections of male and female C57BL/6J mice (2.5 months) across global demyelination and control conditions^31^. Global demyelination models consisted of induction of experimental autoimmune encephalomyelitis (EAE) via injection of myelin oligodendrocyte glycoprotein (MOG). We downloaded processed data from https://zenodo.org/records/8037425 and selected only whole brain coronal sections. We mapped several cell types to our cell type classifications and used existing region annotations. The percentage of zero counts in this dataset (76%) was similar to that in the coronal section dataset used for training the spatial aging clocks (70%).

### MERFISH localized demyelination injury data processing

We applied our spatial aging clocks to publicly available MERFISH spatial transcriptomics data of three whole brain coronal sections at different depths collected from young (3-4 months) male C57BL/6J that were subjected to demyelination injury via stereotactic injection of lysophosphatidylcholine at coordinates (from bregma): (X, ± 1.0 mm; Y, −0.1 mm)^29^. We downloaded processed data and metadata from Gene Expression Omnibus (GSE202638) and mapped several cell types to our cell type classifications. Due to the spatially localized nature of the demyelination injury and the lack of control conditions in this dataset, we analyzed the predicted ages of cells in this dataset by spatially visualizing all cells by their positive age acceleration (negative values floored at zero) across each of the coronal sections to determine spatial patterns in age acceleration with respect to the site of injury for all cell types and for different major cell types. The percentage of zero counts was higher in this dataset (94%) than in the coronal section dataset used for training the spatial aging clocks (70%).

### Effect of interventions on predicted age

For a given cell type and experiment, to quantify the difference in predicted ages between two experimental conditions, we computed the difference in median predicted ages between cells of the two conditions. Specifically for interventions, we subtract the median predicted age of cells belonging to animals in the control condition (sedentary, control) from the median predicted age of cells belonging to animals in the intervention condition (exercise, OSKM, LPS, AD). Positive values indicate that the intervention has an accelerated aging effect and negative values indicate that the intervention has a rejuvenating effect on the cells of the animals. For the whole-body partial reprogramming experiment, since there was a difference in the mean age of animals in the control and OSKM conditions of 0.3 months, we corrected for this difference by adding an intercept to the predicted ages before computing the effect. We computed this effect at the global levels (all cells of a cell type) and at the regional level (all cells of a cell type within a defined anatomic region). For the adverse interventions datasets (LPS and Alzheimer’s disease), we used the existing anatomic region annotations and mapped them to closest region label in our study. For all global comparisons, we quantified the difference in median predicted age and performed two-sided Mann-Whitney U-test on both the predicted ages of all cells and the median predicted ages of all mice between different conditions. These statistics are reported in Supplementary Table 12.

Spatial visualizations of the effect of interventions on predicted age involved computing an effect value for each cell in a sample in the intervention condition. Specifically, for each cell in the intervention condition, we subtract the median predicted age of cells belonging to animals in the age-matched control condition (sedentary, control) from the predicted age of that cell, and then visualize that cell by its spatial coordinates and colored by the computed value.

### Cell proximity effects of aging and rejuvenation

To compute the distribution of cell neighbor to neighbor distances, we constructed a triangulation mesh graph connecting neighboring cells on a given sample using squidpy.gr.spatial_neighbors with delaunay=True^178^. We used the centroid of each cell to compute distances between cells. We computed subregion-specific distance cutoffs for calling nearby cells as the average of the median neighbor-neighbor distances measured across all samples. The distance cutoffs (in microns) for the aging (coronal) study were: CC/ACO: 24.89, CTX_L1/MEN: 25.91, CTX_L2/3: 24.05, CTX_L4/5/6: 27.24, STR_CP/ACB: 21.65, STR_LS/NDB: 20.36, VEN: 17.86. The distance cutoffs (in microns) for the exercise study were: CC/ACO: 23.58, CTX_L1/MEN: 22.13, CTX_L2/3: 21.80, CTX_L4/5/6: 24.81, STR_CP/ACB: 20.75, STR_LS/NDB: 19.82, VEN: 16.23. Using these subregion-specific distance cutoffs, we identified target cells near effector cells (“Near”) and matched them to target cells far from any effector cells (“Far”) for a given target cell type and effector cell type. First, we compute for each cell, the shortest Euclidean distance to any effector cell in the same sample for each effector cell type. Then, for each sample and combination of target cell type and effector cell type, we label all target cells with shortest Euclidean distance to effector cell type that is less than the corresponding distance cutoff as “Near” and match them with “Far” target cells in the same subregion and sample that are farthest away from the effector cell type with shortest distance greater than the cutoff for “Near” cells. After obtaining matched sets of “Near” and “Far” target cells across all samples, we combine them into a single set to estimate the proximity effect of the effector cell type on the target cell type. The proximity effect is computed as Cohen’s *d* measure of effect size between the age acceleration of the “Near” target cells compared to the age acceleration of the “Far” target cells. We filtered out comparisons with less than 50 “Near” cells or less than 50 “Far” cells. Positive proximity effect values indicate a pro-aging effect exerted by the effector cell type on the target cell type and negative proximity effect values indicate a pro-rejuvenating effect exerted by the effector cell type on the target cell type. We also compute the normalized frequency of proximity interactions as the number of “Near” target cells divided by the total number of target cells in the study and the statistics and *P*-value from the associated two-sided Student’s t-test between the age acceleration of the “Near” and “Far” target cells.

We also implemented several variations to the standard proximity effect analysis. For region-specific proximity effects, we restricted the proximity effect analysis to only cells within each of the anatomic regions. For cell proximity effects using alternative definitions of “Far” cells, we used two orthogonal approaches. The first approach consisted of random sampling of “Far” cells from all target cells with shortest distance greater than the cutoff for “Near” cells in the same subregion and sample. The second approach consisted of selecting the set of “Far” cells with total raw transcript counts closest to the mean total raw transcript count of the “Near” cells in the same subregion and sample. For cell proximity effects based on predicted ages obtained from non-spatial aging clocks such as single-cell aging clocks using cell type-specific pseudobulking of gene expression (1000 bootstrap samples of 30 cells following prescribed procedures^3^) during training and prediction [SingleCell (PB)] or spatial aging clocks that did not use the SpatialSmooth step for prediction [SingleCell (SS)], we used predicted ages from these aforementioned clocks in lieu of predicted ages from the spatial aging clocks and performed cell proximity analysis following the original setup. For the “Spillover Filtered” cell proximity effects, we used predicted ages from the spatial aging clocks trained on the 220 genes with less than 5% estimated spillover based on an internal metric using gene expression variations influenced by local cellular composition. For cell proximity effects on datasets other than the coronal section dataset, we computed dataset-specific distance cutoffs for subregions/regions and computed proximity effects using the original setup for all cell types that were mapped to our cell type classifications. We calculated separate proximity effects for cells across the entire external dataset and for cells restricted to the control conditions. To assess the effect of interventions on cell proximity effects, we computed cell proximity effects separately for each experimental condition.

The average proximity effect for each effector cell type was computed as the average proximity effect across all target cell types after filtering. To assess the most impacted target cell types, the mean absolute proximity effect for each target cell type was computed by averaging the absolute value of the proximity effect across all effector cell types after filtering.

### Spatial permutation analysis

We performed global spatial permutations of cells as a negative control for the spatial proximity effects measured for effect cells. In each permutation, for each sample in the aging (coronal) study, we randomly permuted the spatial coordinates of each cell, equivalent to shuffling cell type labels, using numpy.random.permutation. This permutation was performed across the entire sample. Then, we measured proximity effects on the permuted dataset. We repeated these permutations twenty times using random seeds drawn uniformly from 0 to 50,000 with a generating random seed of 444. We computed the median and 95% confidence interval for the proximity effect of each effector cell type across these twenty permutations.

### Area-restricted cell proximity effects

To remove the potential influence of transcript spillover from cell segmentations on the cell proximity effect analysis, we developed an alternative approach to compute area-restricted cell proximity effects, where “Near” cells are labeled using two cutoff distances instead of the single cutoff distance in the standard proximity effect analysis (see Extended Data Fig. 9m). We set the larger of the two cutoff distances equal to the subregion-specific distance cutoff from “<u>Cell proximity effects of aging and rejuvenation”</u>. We set the smaller cutoff distance to be 15 microns less than the larger cutoff distance. We label target cells with shortest Euclidean distance to effector cell type that is greater than the smaller distance cutoff but less than the larger distance cutoff as “Near” and match them with “Far” target cells in the same region and sample that are farthest away from the effector cell type with shortest distance greater than the larger distance cutoff for “Near” cells. Then, area-restricted cell proximity effects are calculated using the same approach as described in “<u>Cell proximity effects of aging and rejuvenation”</u> except with these modified “Near” and “Far” cell labels.

### Spatial visualization of age acceleration near effector cells

We generated visualizations of cells colored by their age acceleration around key effector cell types (e.g. T cell and NSCs) by selecting an effector cell, drawing a square bounding box centered on the effector cell with edge lengths of 200 microns, and then visualizing all cells with center coordinates inside of the bounding box. We manually selected visualizations from a middle-aged mouse (18.8 months) for T cells and NSCs.

### Spatial gradation of cell proximity effects

We computed cell proximity effects as a function of the unit distance between the effector and target cells, where the unit distance is defined as a scalar value that is multiplied by the subregion-specific distance cutoffs from “<u>Cell proximity effects of aging and rejuvenation”</u>. These unit distances are used as new cutoff distances to compute cell proximity effects. We also analyzed spatial gradation of area-restricted cell proximity effects by setting the larger of the two cutoff distances to the unit distance and the smaller cutoff distance to be 15 microns less than the larger cutoff distance, which ensured no overlap of cells defined as “Near” between integer multiples of the cutoff distances.

### Activation/inflammation glia signatures and subtype identification

We computed microglia activation scores and oligodendrocyte/OPC inflammation scores by summing the log-normalized expression of all shared genes between gene signature sets curated from several published sets^22,179,180^ and the 300 genes in the MERFISH dataset. The microglia activation score consisted of gene expression from *Apoe, B2m, C1qa, Cd69, Cd9, Il1a, Il1b, Il6, Lyz2*. The oligodendrocyte/OPC inflammation score consisted of gene expression from *C4b, Cdkn1a, H2-D1, Ifit1, Stat1*.

To control for activation/inflammation status in the T cell proximity effect, we selected the cutoff for classifying microglia (activated) and oligodendrocyte (inflammation) subtypes as the top 0.2% highest scores since that was the highest percentage cutoff for which there was no statistically significant difference in the distribution of activation/inflammation scores between activated/inflammation cells that were near or far from T cells. We performed all associated proximity effect analysis using the normal and activated/inflammation subtype labels for microglia and oligodendrocytes.

### Cell type perturbation modeling with deep learning (graph neural networks)

Deep learning approaches such as graph neural networks can be leveraged to predict the effects of *in silico* perturbations. For each section in the coronal section dataset, we constructed a global graph connecting neighboring cells on a given sample using squidpy.gr.spatial_neighbors with delaunay=True^178^ and pruned edges connecting neighboring cells with distance greater than 200 microns. To define local cell graphs, we extracted induced 2-hop subgraphs of the global graph by random sampling of at most 100 center cells per cell type for T cells and NSCs. To increase the heterogeneity of these local cell graphs, we further augmented these local cell graphs by inducing 2-hop subgraphs centered on all cells within the first set of local subgraphs. For each local cell graph, we defined the neighborhood aging as the average age acceleration of all cells in the graph. We defined cell (node) features as a one-hot vector for cell type using the 18 cell type annotations. We trained a graph neural network (GNN) model using PyTorch Geometric^181^ to predict neighborhood aging from the features of a local cell graph. The GNN model consisted of a three-layer graph isomorphism network (GIN), with node embedding updates for each layer modeled by a linear transformation with hidden dimensionality of 16 followed by batch normalization, ReLU transformation, and a final linear transformation with hidden dimensionality of 16. The first two GIN layers were followed by a ReLU transformation and the output of the final GIN layer was globally pooled by addition before linearly transformed to predict neighborhood aging. We trained the GNN model using a balanced mean-squared error loss^182^, the Adam optimizer^183^ with a learning rate of 0.0001, and a batch size of 64 for 50 epochs.

To model “loss of function” perturbations for a given local cell graph, we mutated the center cell of the graph into a random cell type drawn from a uniform distribution across all cell types in the dataset excluding the original cell type. We performed “loss of function” perturbations for local cell graphs that had T cells and NSCs as center cells. To model “gain of function” perturbations for a given local cell graph, we mutated the center cell of the graph into the specified cell type by modifying its node features. We performed “gain of function” perturbations for all local cell graphs and mutated the center cells to either T cells or NSCs. We excluded “gain of function” perturbations resulting in unperturbed local cell graphs from our analysis (e.g. graph with T cell as center cell mutated to T cell). For both “loss of function” and “gain of function” experiments, we used endothelial cells as a negative control setting. Using the GNN model, we evaluated the effect of both “loss of function” and “gain of function” perturbations by predicting the neighborhood aging for the unperturbed local cell graph and the neighborhood aging for the perturbed local cell graph. The effect on neighborhood aging is then measured as the unperturbed neighborhood aging subtracted from the perturbed neighborhood aging (positive values indicating a pro-aging perturbation and negative values indicating a pro-rejuvenating perturbation).

### TISSUE imputation

We filtered cells in the aging (coronal) study to include NSCs, neuroblasts, T cells, and all target cells labeled “near” or “far” with respect to NSCs or T cells under the proximity effect framework. We re-centered each sample to grid lattice points such that no samples spatially overlap. In addition to “near” and “far” labels, we also labeled each T cell and NSC by the strength of their proximity effects. First, we compute the neighborhood age acceleration as the average age acceleration of all cells within the maximum subregion distance cutoff for each T cell or NSC. Then, we labeled all T cells with the 50% highest neighborhood age acceleration as “strong” and the remaining T cells as “weak”, and we labeled all NSCs with the 50% lowest neighborhood age acceleration as “strong” and the remaining NSCs as “weak”.

TISSUE is an algorithm that provides an uncertainty-aware framework for performing differential gene expression/signature analysis on imputed spatial gene expression with marked reductions in false discovery rates compared to other approaches^40^. We applied the TISSUE algorithm on the filtered MERFISH data for whole-transcriptome uncertainty-aware spatial gene expression imputation using dissociated single-cell RNAseq datasets of the mouse striatum, ventricle, and corpus callosum across multiple ages^2,3^. Log-normalized gene expression data from the first dissociated single-cell RNAseq dataset^2^ was used for imputation of gene expression for T cells (8,170 genes imputed in total), and log-normalized gene expression data from the second dissociated single-cell RNAseq dataset^3^ was used for imputation of gene expression for all other cell types (12,719 genes imputed in total). We used the raw MERFISH counts normalized by cell segmentation volume as input for imputation. We used TISSUE with the SpaGE imputation algorithm^76^ and Tangram imputation algorithm^77^, under default settings such as 10 folds of cross-validation, 4 stratified gene groups, 1 stratified cell groups, and 100 multiple imputations (for hypothesis testing purposes). Evaluation of TISSUE calibration quality and SpaGE/Tangram imputation performance were conducted using the TISSUE software package and associated code^40^.

### TISSUE differential gene expression analysis and pathway enrichment

We used TISSUE to perform uncertainty-aware differential gene expression analysis on the imputed gene expression. We performed two sets of comparisons using TISSUE hypothesis testing of gene signatures. First, we compared gene signature scores between “near” and “far” target cells with respect to either T cells or NSCs. Second, we compared gene signature scores between “strong” and “weak” T cells or NSCs. We determined significantly differentially expressed genes (DEGs) as genes with permissive cutoff of *P* < 0.05 from TISSUE two-sample t-tests for both SpaGE and Tangram imputed expression to reduce false discoveries resulting from variability or biases in imputation quality. We further filtered these genes such that the associated TISSUE t-statistic were of the same sign across the tests on SpaGE and Tangram imputations. We did not perform any filtering of genes by the log fold change in imputed gene expression across conditions because unlike the TISSUE *P*-value, the log fold change is not calibrated for uncertainty in spatial gene imputation.

GO biological process (2023) gene set enrichment was performed using the EnrichR framework^184^ accessed through gseapy^185^ (version 1.0.4). We separated DEGs for each comparison into upregulated DEGs (positive TISSUE t-statistic) and downregulated DEGs (negative TISSUE t-statistic) and used these gene lists as inputs into EnrichR to obtain enrichment statistics for different gene sets. We selected the top 5 gene sets for each comparison, which were generally representative of all significantly associated gene sets.

### Test for equal variances between groups

We performed Levene’s test for equal variances using the scipy.stats.levene implementation with default settings (center=median, proportiontocut=0.05) for analyses relying on Cohen’s *d* or variations of the Student’s t-test. Levene’s test was applied to test for equal variance of age acceleration of “Near” and “Far” cell groupings for each cell proximity effect. Levene’s test was also applied to test for equal variance of TISSUE (SpaGE) imputed gene expression of “Near” and “Far” cell groupings with respect to T cells and NSCs. In both cases, the number of samples between the compared groups was equal, a setting in which Cohen’s *d* and Student’s t-test are usually robust against unequal variances across groups.

### TISSUE gene signature scores

We further modified the hypothesis testing framework in TISSUE to perform testing for gene signature scores (sum of imputed gene expression values across all genes in a GO gene set). We have made this modification publicly available in the TISSUE package^40^ as tissue.downstream.multiple_imputation_gene_signature and we set n_imputations=100 for all gene signature tests. For imputed gene signature scores, we performed the same two sets of comparisons as outlined in “TISSUE differential gene expression analysis and pathway enrichment”. For similar comparisons involving expression of individual genes (e.g. *Bst2*, *Stat1*, *Cd9*, *Vegfa*) instead of gene signatures, we used the measured (log-normalized) gene expression values in the MERFISH dataset instead of imputed values.

### Immunofluorescence staining of brain sections

All immunostainings were performed on male C57BL/6JN mice at the indicated ages. Mice were sedated with 0.8 mL 2.5% vol/vol Avertin (Sigma-Aldrich, T48402-25G) in PBS (Corning, 21-040-CV) and perfused via the left ventricle of the heart with 5 mL of PBS (Corning, 21-040-CV) with heparin sodium salt (50 U ml, Sigma-Aldrich, H3149-50KU) to remove circulating blood cells followed by 4% PFA (Electron Microscopy Sciences, 15714) in PBS. Brains were post-fixed overnight in 4% PFA (Electron Microscopy Sciences, 15714) and then dehydrated in 30% sucrose (Sigma-Aldrich, S3929) for 72 hours. Brains were embedded in O.C.T. compound (Fisher Healthcare Tissue Plus, 4585), sectioned in 20-μm coronal sections using a cryostat (Leica, CM3050S), and then mounted on electrostatic glass slides (Fisher Scientific, 12-550-15). Coronal sections at a similar depth to the MERFISH coronal sections were collected. For the following steps, all brain sections were processed simultaneously within each experiment. For immunofluorescence staining, brain sections were washed with PBS for 5 minutes, permeabilized in ice-cold methanol with 0.1% Triton X-100 (Fisher Scientific, BP151) for 15 minutes, and washed 3X with PBS for 5 minutes. We performed antigen retrieval by placing the brain sections in 10mM sodium citrate buffer (pH 6.0; 2.94 g Tri-sodium citrate dihydrate (Sigma-Aldrich, S1804) in 1,000 ml milliQ H2O adjusted to pH 6.0 with 1 N HCl) + 0.05% TWEEN 20 (Sigma-Aldrich, P1379-1L) at 85°C in a water bath for 2 hours. Brain sections in buffer were cooled to room temperature for 20 minutes and then washed 2X with PBS for 3 minutes. Sections were blocked for 30 minutes at room temperature with block buffer consisting of 5% normal donkey serum (ImmunoReagents, SP-072-VX10) and 1% BSA (Sigma, A7979) in PBS. Primary antibodies were diluted in blocking buffer and primary antibody staining was performed overnight at 4°C. Primary antibodies used were as follows: anti-GFAP (1:1000 dilution, Abcam, ab53554), anti-Ki67 (1:500 dilution, Thermo Fisher Scientific, 14-5698-82), anti-DCX (1:500 dilution, Millipore Sigma, AB2253), anti-EGFR (1:200 dilution, Millipore Sigma, 06-847), anti-STAT1 (1:500 dilution, Cell Signaling Technology, 14994), anti-CD3 (1:500, Abcam, ab11089), anti-S100A6 (1:500, Abcam, ab181975), anti-CD9 (1:100, Thermo Fisher, eBioKMC8 (KMC8)), anti-CPT1A (1:200, Abcam, ab128568). Sections were washed 3X with PBS and 0.2% TWEEN 20 for 10 minutes at room temperature followed by 2X with PBS for 15 minutes. Secondary antibodies were diluted in blocking buffer and secondary antibody staining was performed for 2 hours at room temperature. Secondary antibodies used were as follows: Donkey anti-Goat 647 (1:500 dilution, Invitrogen, A21447), Donkey anti-Rabbit 488 (1:500 dilution, Invitrogen, A-21206), Donkey anti-Guinea pig 594 (1:500 dilution, Jackson ImmunoResearch, 706-585-148), Donkey anti-Rat 647 (1:500 dilution, Invitrogen, A48272), Donkey anti-Rat 488 (1:500, Invitrogen, A21208), Donkey anti-Mouse 647 (1:500, Invitrogen, A31571), Donkey anti-Rabbit 568 (1:500, Invitrogen, A10042). DAPI (1:500, Thermo Fisher, 62248) was added during secondary antibody staining. Sections were washed 3X with PBS and 0.2% TWEEN 20 for 10 minutes followed by 3X with PBS for 5 minutes. Sections were mounted with ProLong Gold Antifade Mountant with DAPI (Thermo Fisher, P36931) and a coverslip.

For immunofluorescence experiments corresponding to spatiotemporal marker expression, images were acquired on a Zeiss LSM 900 with Zeiss ZEN Blue 3.0 software or a Zeiss LSM 980 confocal microscope with Zeiss ZEN Blue 3.3 software using a 10x objective and automatic tiling of entire coronal brain sections. The same equipment and microscope acquisition settings were used for different brain sections with the same antibody panel. Tile images were stitched using Zeiss ZEN Blue software. Image brightness and contrast were adjusted in ImageJ (1.53n) to enhance visualization with the same settings applied to all images shown for each antibody staining panel.

For immunofluorescence experiments corresponding to T cell and NSC proximity effects, images were acquired with a 60x objective on a Nikon Eclipse Ti confocal microscope equipped with a Zyla sCMOS camera (Andor) and NIS-Elements software (AR 4.30.02, 64-bit). For immunofluorescence experiment corresponding to T cell proximity effects, we acquired at least two images containing T cells and at least one image without any T cells from each of four anatomic regions (corpus callosum (CC), cortex (CTX), striatum and adjacent regions (STR), and lateral ventricles (VEN)) for each brain section. The same image acquisition settings were used for all brain sections in this experiment. For immunofluorescence experiment corresponding to NSC proximity effects, we acquired individual images tiling the entire right and left lateral ventricles for each brain section. The same image acquisition settings were used for all brain sections in this experiment.

### Immunofluorescence quantification of T cell proximity effect mediated through interferon response

Immunofluorescence imaging was performed on brain sections from old (28 months) mice. All cell segmentations and cell type annotations were performed using automated pipelines in QuPath 0.5.1. For all images, cell nuclei were automatically segmented based on DAPI intensity, and then nuclear segmentation masks were expanded by 2 μm to define the cell segmentations. Cells were labeled as CD3^+^ using a manually determined threshold for mean cell CD3 intensity. The same threshold was used across all images and CD3^+^ cells were annotated as T cells. For all CD3^-^ cells, we defined cells as “Near” if they were located within 50 μm of a T cell based on the centroids of cell segmentations and otherwise defined them as “Not Near”. For each anatomic region (CC, CTX, STR, VEN) and each cell proximity definition (“Near” or “Not Near”), we quantified the mean STAT1 intensity by averaging the mean cell STAT1 intensity across each section and then by averaging these values across each mouse (3-5 coronal sections per mouse, 8 mice per condition). The mean STAT1 intensity values were normalized for each independent experiment by dividing them by the corresponding mean “Not Near” STAT1 intensity.

### Immunofluorescence quantification of NSC proximity effect mediation through exosomes and fatty acid oxidation

Immunofluorescence imaging was performed on brain sections from young (3.5 months) mice. All cell segmentations and cell type annotations were performed using automated pipelines in QuPath 0.5.1. For all images, regions of interest were defined along the lining of the lateral ventricles. Within these regions of interest, cell nuclei were segmented based on DAPI intensity, and then nuclear segmentation masks were expanded by 2 μm to define the cell segmentations. Cells were labeled as S100A6^+^ using a manually determined threshold for mean nuclear S100A6 intensity. The same threshold was used across all images and S100A6^+^ cells were annotated as NSCs and other cells were annotated as non-NSCs. To compare CD9 intensity between S100A6^+^ and S100A6^-^ cells, we quantified the mean CD9 intensity by averaging the mean cell CD9 intensity across each section and then by averaging these values across each mouse (3 coronal sections per mouse, 5 mice per condition). The mean CD9 intensity values were normalized for each independent experiment by dividing them by the CD9 intensity in S100A6^-^ cells. For all cells, we defined cells as “Near” if they were located within 20 μm of a NSC (excluding itself) based on the centroids of cell segmentations and otherwise defined them as “Not Near”. To compare CPT1A intensity between “Near” and “Not Near” cells, we quantified the mean CPT1A intensity by averaging the mean cell CPT1A intensity across each section and then by averaging these values across each mouse (3 coronal sections per mouse, 5 mice per condition). The mean CPT1A intensity values were normalized for each independent experiment by dividing them by the corresponding mean “Not Near” CPT1A intensity. To examine the correlation between CPT1A intensity of all “Near” cells and the CD9 intensity of the nearest NSC, we matched all “Near” cells to their nearest NSC defined by the minimum non-zero Euclidean distance between centroids and computed the Pearson and Spearman correlation between the paired CPT1A and CD9 intensities. To check the robustness of the correlation analysis to technical variability, we also re-imaged all sections using a Zeiss LSM 900 with Zeiss ZEN Blue 3.0 software and performed the same image analysis. Results were highly consistent with the reported findings.

### Statistics and Reproducibility

For measures of Pearson and Spearman correlation and their associated test statistics (*P*-values and 95% confidence intervals), we used the implementation from scipy.stats.pearsonr and scipy.stats.spearmanr. For most comparisons, we used the two-sided Mann-Whitney tests according to the implementation from scipy.stats.mannwhitneyu. We used Cohen’s *d* for cell proximity effect analysis in which similar variances in age acceleration were typically observed between “Near” and “Far” cell groups (Extended Data Fig. 9a). GO biological process enrichment analysis was performed with topGO^176^ (R package version 2.54.0) using Fisher’s exact test for all Biological Process terms and using all other genes measured by MERFISH as background.

For the gene signature and differential gene expression analysis with imputed spatial gene expression, we used the two-sample two-sided TISSUE t-test with specifications outlined in the “TISSUE imputation” section. The TISSUE t-test is the only statistical test that considers uncertainty in imputed gene expression and to further decrease false discoveries, we selected genes with consistent differential expression across two imputation methods (SpaGE^76^ and Tangram^77^). For most comparisons using the TISSUE t-test, variances in imputed gene expression were similar across groups (Extended Data Fig. 10a). We did not perform multiple hypothesis correction of *P*-values for the targeted gene signature analysis using TISSUE imputation because gene signatures were manually selected. Reported *P*-values are from the SpaGE imputed gene expression. For pathway enrichment for differentially expressed genes identified with TISSUE and imputed gene expression, we used gseapy.enrichr with the ‘GO_Biological_Process_2023’ gene set, ‘mouse’ as the organism, and all imputed genes as background.

Technical reproducibility of MERFISH measurements was confirmed by profiling consecutive sections in the 140-gene MERFISH dataset^40^ (Extended Data Fig. 1a). For the MERFISH (300 gene) aging dataset, we combined two cohorts of animals (n=10 for each cohort) and confirmed reproducibility of spatial aging clock performance and generalization of key findings within each cohort (Extended Data Fig. 5b,c and Extended Data Fig. 9g,h). All experimental validation was performed on independent mice.

---

**Extended Data Figure 1:**
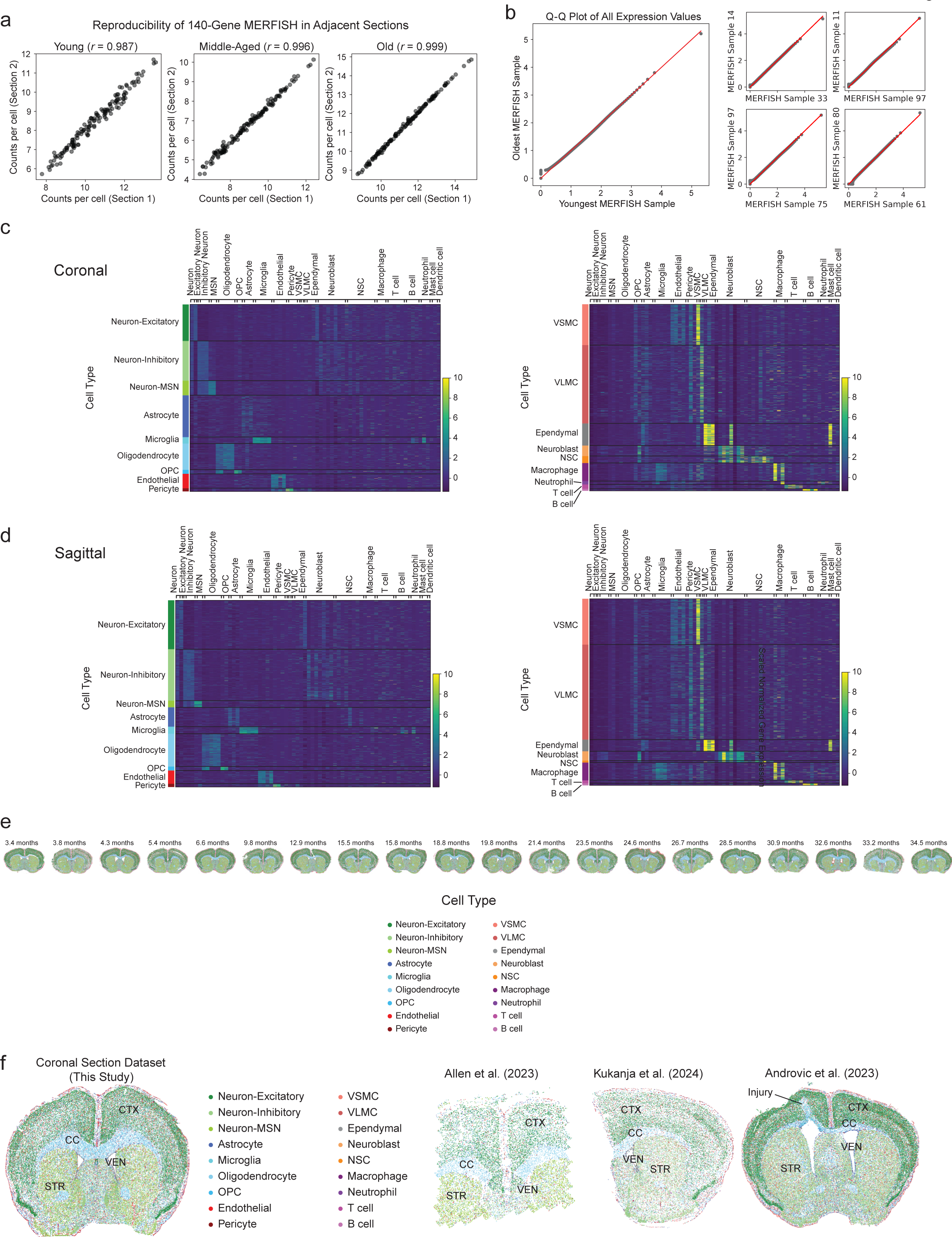
MERFISH data reproducibility and external validation. **a,** Reproducibility plot of individual genes by their counts per cell between two adjacent coronal sections in the 140-gene MERFISH dataset. Pearson correlation between the gene counts per cell of the two sections is noted in parentheses. **b,** Q-Q (quantile-quantile) plot of all log-normalized expression values across all cells and genes in the MERFISH data of the youngest mouse (3.4 months) compared to the oldest mouse (34.5 months) in the coronal sections dataset (left) and for four randomly selected pairs of mice from the coronal sections dataset (right). **c,d,** Heatmaps showing the scaled log-normalized expression of key cell type marker genes for different cell types (columns) and grouped by the identified cell type clusters (rows) for **(c)** the coronal sections of the aging study and **(d)** the sagittal sections of the aging study. **e,** Scatter plot of cells by their spatial coordinates across all coronal sections and ages with cells colored by cell type. **f,** Scatter plot of cells by their spatial coordinates with cells colored by cell type for representative coronal sections from our coronal section dataset (leftmost) and each of three publicly available spatial transcriptomics datasets of adult mouse brain (coronal brain sections). Regions are labeled for each dataset: cortex (CTX), striatum and adjacent regions (STR), corpus callosum (CC), lateral ventricle (VEN), and injury site (arrow).

**Extended Data Figure 2:**
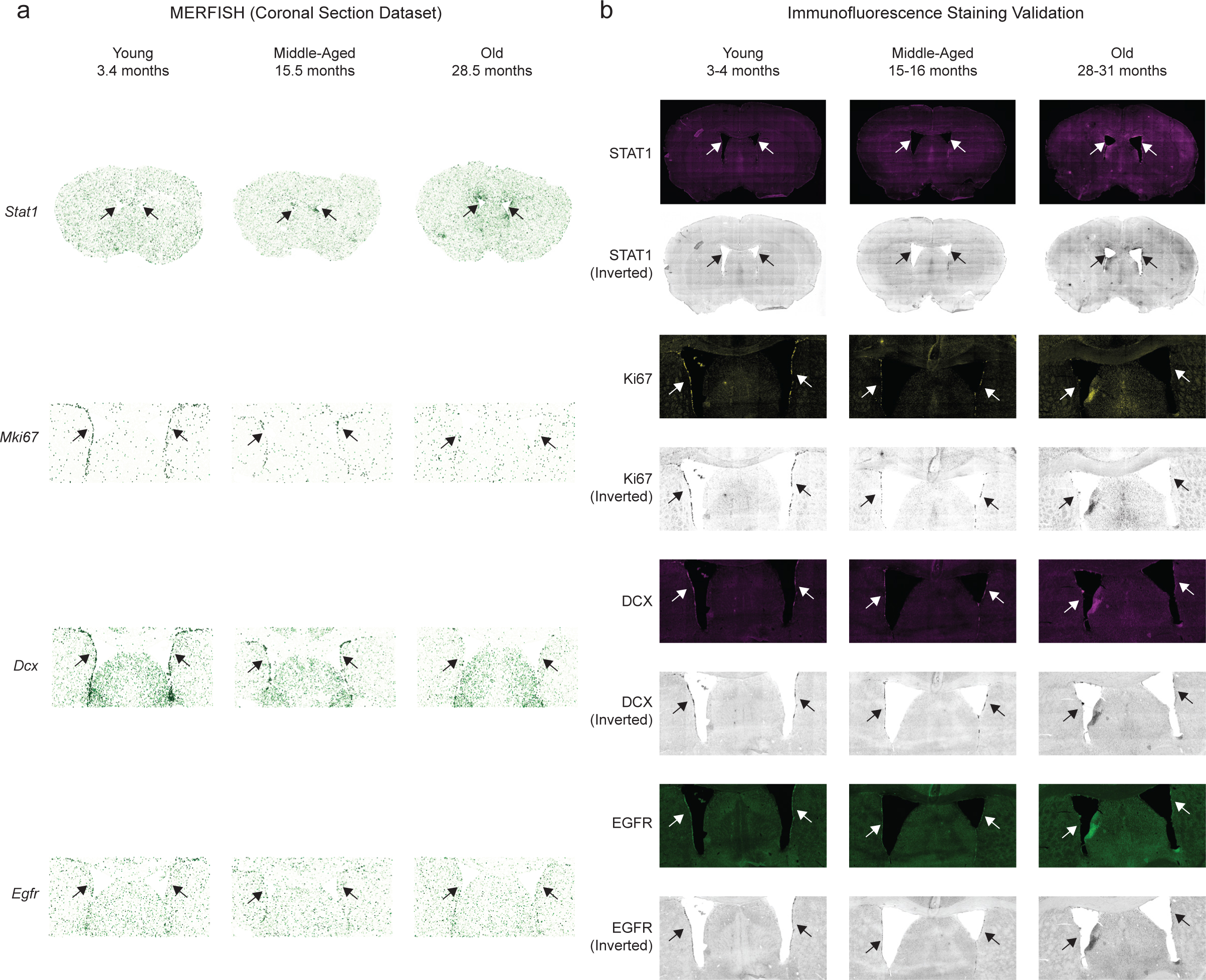
Validation of spatiotemporal gene expression in the MERFISH panel by immunofluorescence staining. **a**, MERFISH coronal section dataset: Scatter plot of cells by their spatial coordinates across all sections for three ages (young 3.4 months, middle-aged 15.8 months, old 28.5 months) with cells colored by scaled log-normalized gene expression of key marker genes in the MERFISH coronal section dataset for 4 genes (*Stat1*, *Mki67*, *Dcx*, *Egfr*). Stat1 is an interferon-response gene. *Mki67* is a cell proliferation marker gene. *Dcx* is a neuroblast marker gene. *Egfr* is a neural stem cell marker gene. The whole coronal sections are shown for *Stat1* and are reproduced from Fig. 1j. The areas around the lateral ventricles are shown for *Mki67*, *Dcx*, and *Egfr*. Arrows indicate the subventricular zone neurogenic niche in the lateral ventricles. **b,** Immunofluorescence validation: Immunofluorescence staining of perfused mouse brain sections from male mice at three ages (young 3-4 months, middle-aged 15-16 months, old 28-31 months) for protein markers corresponding to each of the select markers. The whole coronal section is shown for STAT1. The area around the lateral ventricles is shown for Ki67, DCX, and EGFR. Arrows indicate the subventricular zone neurogenic niche in the lateral ventricles. Shown are the original immunofluorescence images and the inverted grayscale images (to provide a more direct comparison with the MERFISH data).

**Extended Data Figure 3:**
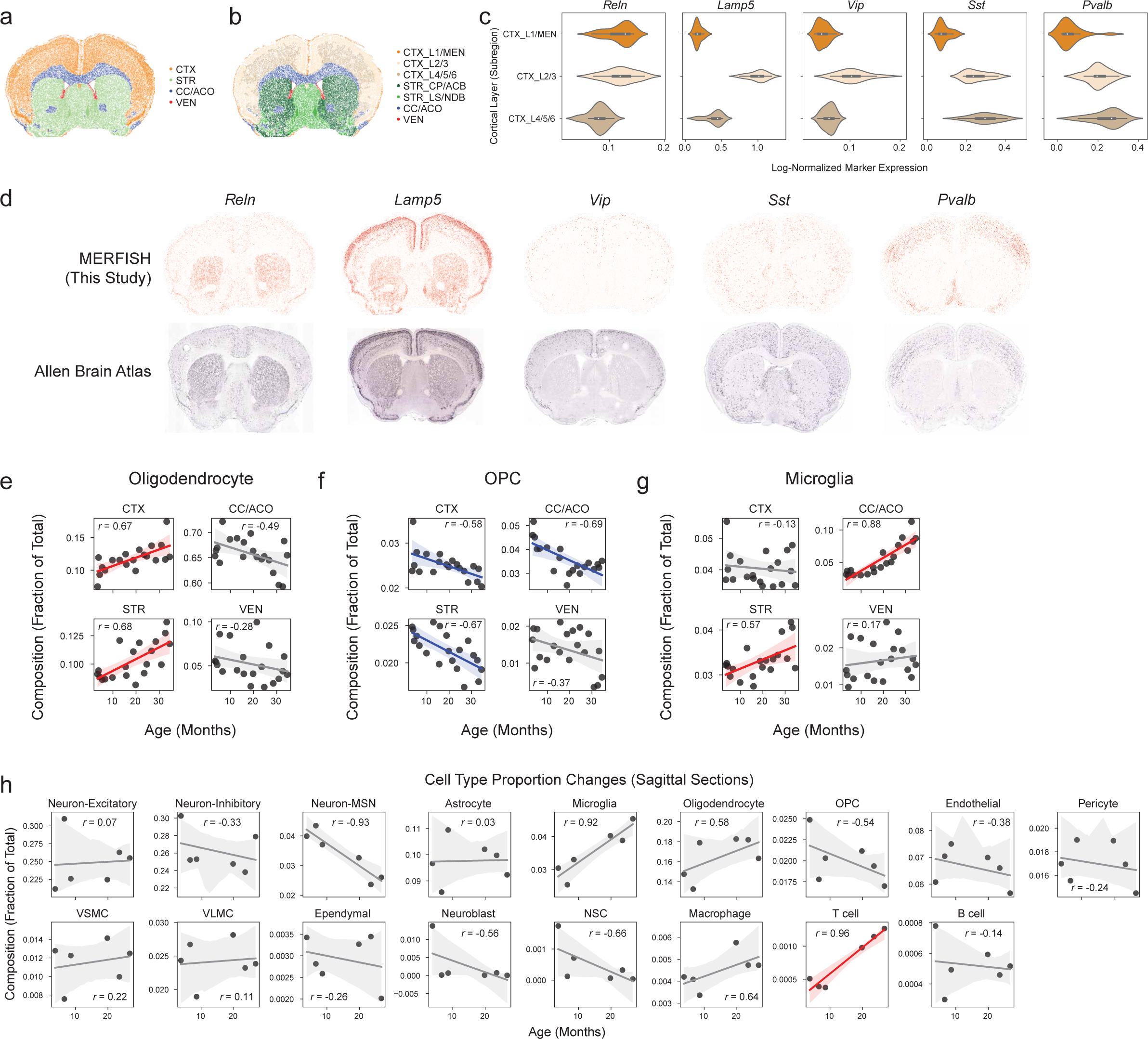
Subregion annotation and cell type composition changes. **a,** Scatter plot of cells by their spatial coordinates in an example coronal section with cells colored by region. **b,** Scatter plot of cells by their spatial coordinates in the example coronal section with cells colored by subregion. **c,** Violin plots showing log-normalized gene expression of five different cortical layer or neuronal subtype markers across all cells in different subregions of the cortex in the coronal section dataset. The markers include: *Reln* (indicative of L1), *Lamp5* (indicative of L2/3 together with L1), *Vip* (indicative of L2/3), *Sst* (indicative of L5 and L6 together with L2/3), and *Pvalb* (indicative of L5 and L6 together with L2/3). The inner box corresponds to quartiles and the whiskers span up to 1.5 times the interquartile range of log-normalized gene expression values. **d**, Scatter plot of cells by their spatial coordinates in the example coronal section with cells colored by their log-normalized expression (upper row) compared to mouse brain coronal sections from the Allen brain in situ hybridization atlas at https://mouse.brain-map.org/ (credit: Allen Institute) (bottom row) for five different cortical layer or neuronal markers. **e-g,** Region-specific composition changes in the coronal section dataset across the four regions (cortex, striatum and adjacent regions, corpus callosum and anterior commissure, lateral ventricles). Line and shaded region correspond to regression line of best fit and corresponding 95% confidence interval. Pearson correlations between cell type composition and age are reported. Significant increases in cell type proportion with age are in red and significant decreases in cell type proportion with age are in blue (Bonferroni corrected *P*-value < 0.05). Shown are plots for **(e)** oligodendrocytes, **(f)** oligodendrocyte progenitor cells (OPCs), and **(g)** microglia. **h,** Global cell type composition changes with each dot representing an individual mouse in the sagittal section dataset with the same plotting and statistical parameters as in (e-g).

**Extended Data Figure 4:**
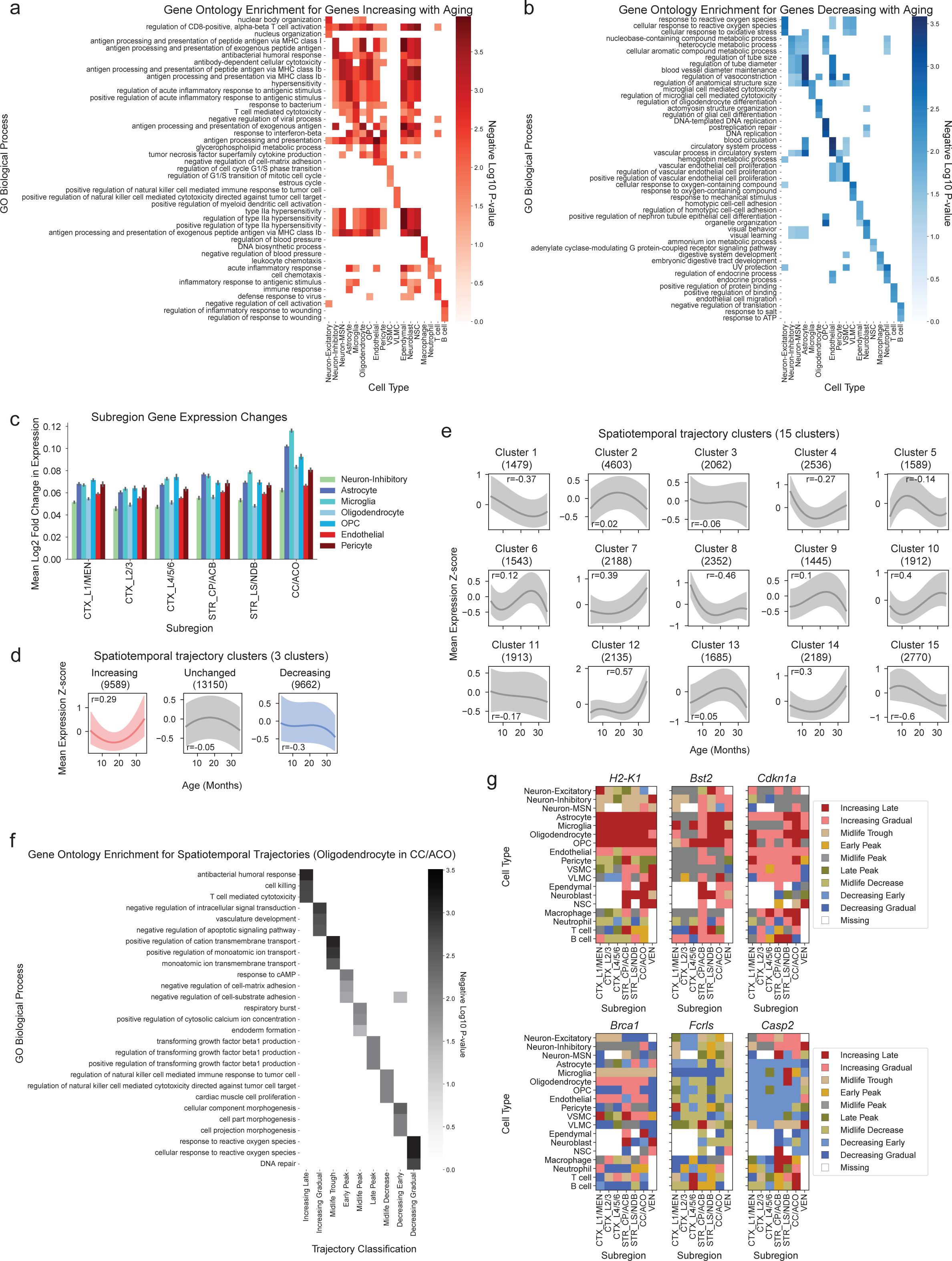
Gene expression changes and trajectories with age. **a,b,** Heatmaps of GO biological process terms with color scale corresponding to negative log10 *P*-value from Fisher’s exact test for **(a)** the top three terms that are significantly enriched for MERFISH genes increasing with age per cell type and **(b)** the top three terms that are significantly enriched for MERFISH genes decreasing with age per cell type. See Supplementary Table 8 for all GO biological process enrichment results. **c,** Mean absolute log2 fold change in gene expression between the youngest five animals and oldest five animals for all cell types with at least 20 cells present in each subregion of each sample. The lateral ventricles (VEN) subregion was excluded due to low numbers of many cell types. Shown are the average values and error bars corresponding to 95% confidence interval for 20 random samples of 20 cells from each cell type/subregion/sample. **d,e**, Median of the mean gene expression z-scores across age of all gene, region, cell type combinations (coronal sections) split into *k* clusters (determined by *k*-means clustering) for **(d)** *k*=3 and **(e)** *k*=15. Shaded regions correspond to interquartile range in expression. All trajectories were smoothed using interpolating B-splines. Pearson correlation (*r*) between median gene expression z-score and age is shown, and the number of trajectories within each cluster is noted inside parentheses. **f,** Heatmap of GO biological process terms with color scale corresponding to negative log10 *P*-value from Fisher’s exact test for the top three terms that are significantly enriched for MERFISH genes in each of the nine annotated gene trajectory clusters for oligodendrocytes in the corpus callosum and anterior commissure region, which corresponds to an abundant cell type in the region with the greatest gene expression change with age (c). See Supplementary Table 9 for all GO biological process enrichment results. **g,** Heatmaps showing spatiotemporal trajectory cluster membership across all subregions and cell types for select genes with distinct patterns: major histocompatibility complex protein-encoding gene *H2-K1* (cell type-specific “Increasing Late”), interferon-response gene *Bst2* (mixed “Increasing Late” and “Increasing Gradual”), cell senescence marker *Cdkn1a* (broad “Increasing Gradual”), DNA damage repair gene *Brca1* (broad “Decreasing Gradual” with “Increasing Gradual” in some cell types), Fc receptor gene *Fcrls* (cell type-specific “Decreasing Gradual”), and cellular apoptosis gene *Casp2* (cell type-specific “Decreasing Early” and “Increasing Gradual”).

**Extended Data Figure 5:**
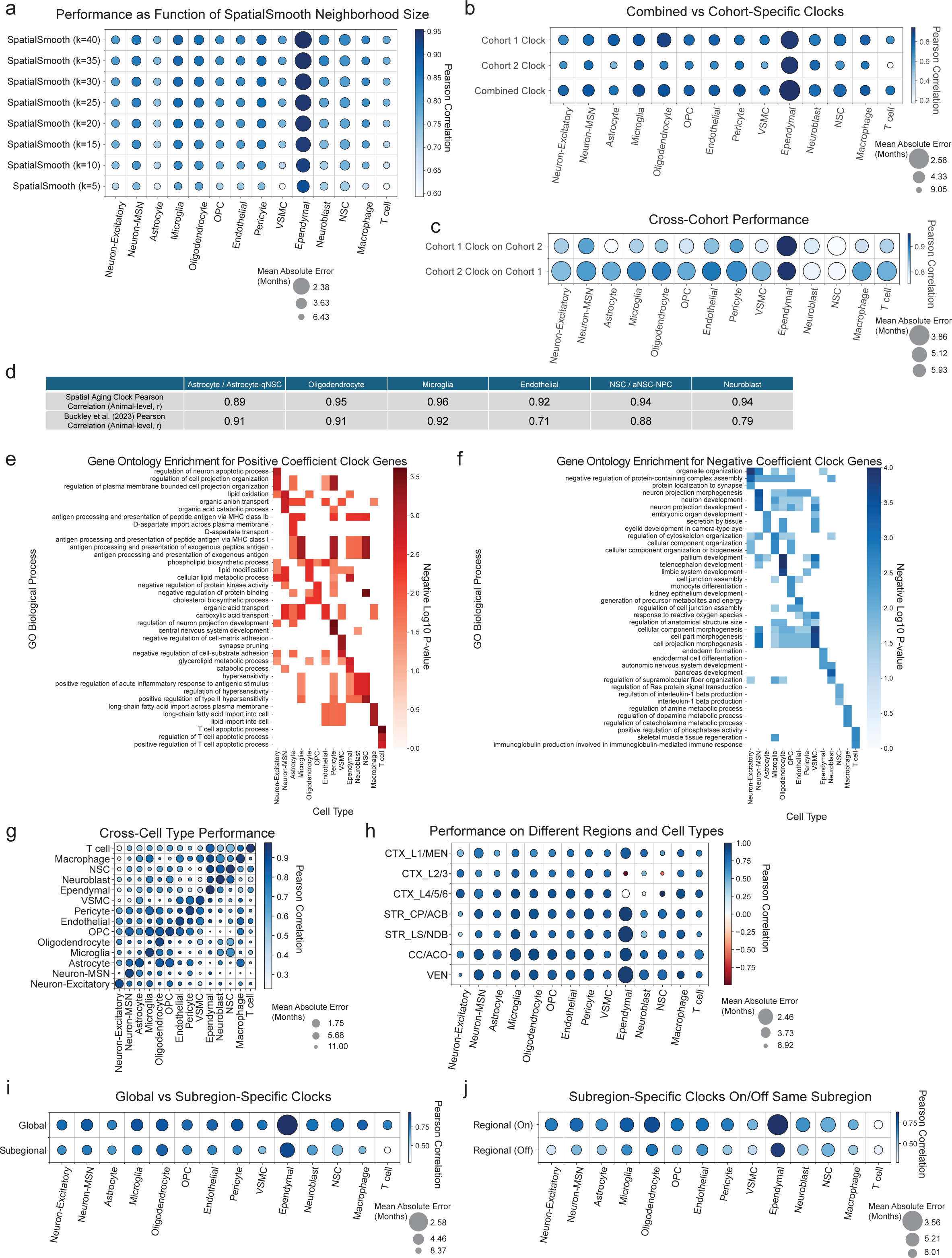
Specificity and robustness of spatial aging clocks. **a-c**, Dot plots evaluating the predictive performance of aging clocks in various contexts. Colors correspond to Pearson correlation between predicted age and actual age and the size of the dots are inversely related to the mean absolute error between predicted age and actual age. Shown are comparisons for **(a)** spatial aging clocks trained and evaluated using different numbers of nearest neighbors in the SpatialSmooth processing step; **(b)** spatial aging clocks trained and evaluated on the two independent cohorts (*n* = 10 per cohort) in the coronal section dataset compared to the spatial aging clock trained on the full (combined) dataset; **(c)** spatial aging clocks trained on one of the independent cohorts (*n* = 10 per cohort) in the coronal section dataset and evaluated on the other independent cohort. **d,** Table comparing Pearson correlation of median predicted age and actual age of individual mice between the spatial aging clocks and existing single-cell RNAseq aging clocks for six shared cell types. Cross-validation was used to generate predicted ages for both clocks within their respective training datasets. **e,f,** Heatmaps of GO biological process terms with color scale corresponding to negative log10 *P*-value from Fisher’s exact test for **(e)** the top three terms that are significantly enriched for genes with positive coefficients per cell type-specific spatial aging clock and **(f)** the top three terms that are significantly enriched for negative coefficients per cell type-specific spatial aging clock. **g-j,** Dot plots evaluating the predictive performance of aging clocks in various contexts. Colors correspond to Pearson correlation between predicted age and actual age and the size of the dots are inversely related to the mean absolute error between predicted age and actual age. Shown are comparisons for **(g)** cell type-specific spatial aging clocks applied to predict age for all other cell types in the coronal aging dataset; **(h)** spatial aging clocks evaluated within each of the subregions in the coronal aging dataset; **(i)** spatial aging clocks (Global) and subregion-specific spatial single-cell aging clocks (Regional) applied to all cells in the coronal aging dataset; **(j)** subregion-specific aging clocks applied to all cells within the same subregion (On) compared to all cells in other subregions (Off).

**Extended Data Figure 6:**
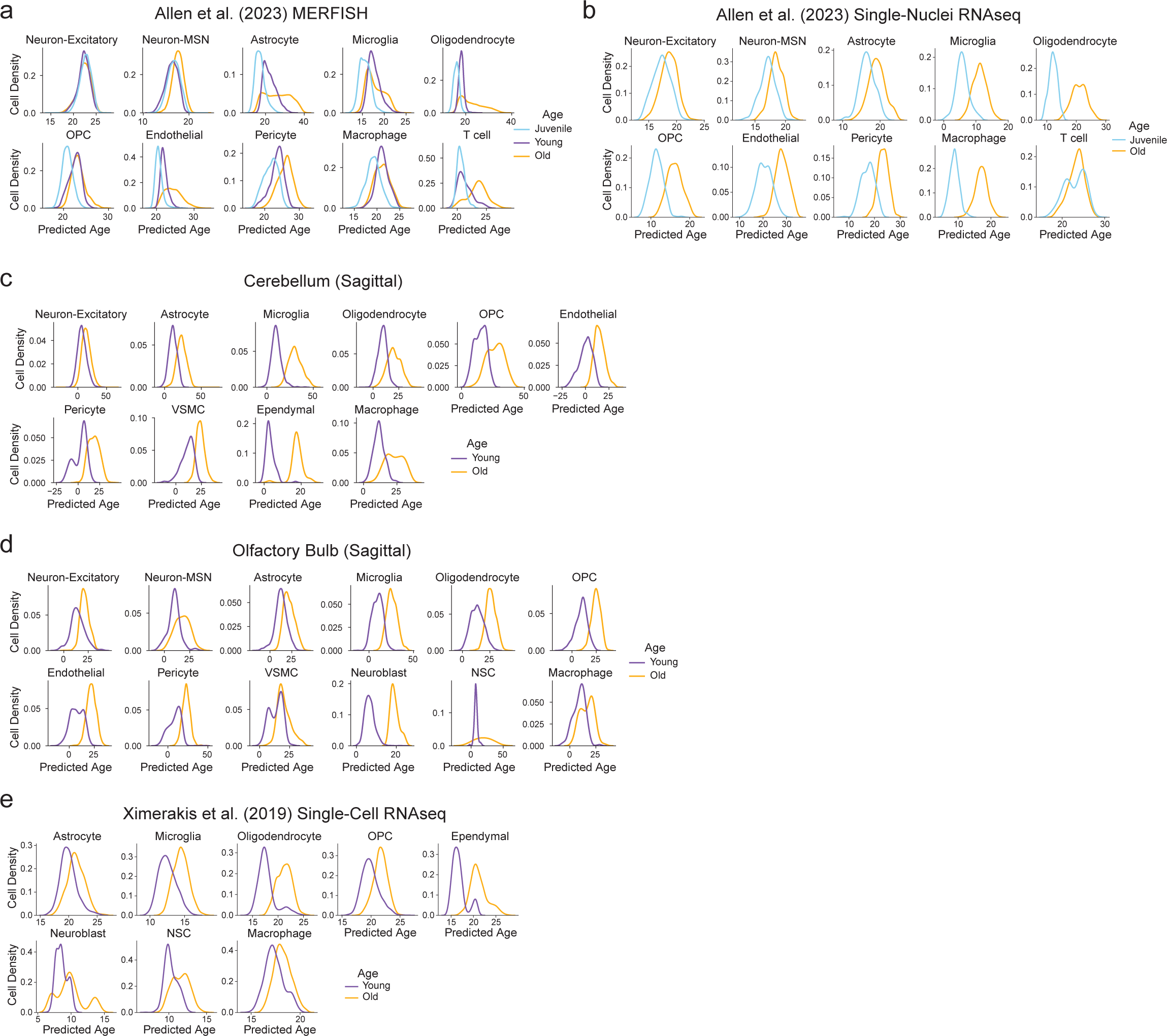
Generalizability of spatial aging clocks. **a-e,** Density of predicted ages computed using spatial aging clocks for different age groups in the following datasets: **(a)** publicly available MERFISH dataset in female mice across juvenile (0.93 months), young (5.58 months), and old (20.93 months) age timepoints, with the dataset sharing 75 genes in common with the coronal section dataset; **(b)** publicly available single-nuclei RNAseq dataset in female mice across juvenile (0.93 months) and old age (20.93 months) timepoints, with the dataset sharing 295 genes in common with the original dataset; **(c)** cerebellum region in the MERFISH sagittal section dataset with young (<9 months) and old (>19 months) age bins; **(d)** olfactory bulb region in the MERFISH sagittal section dataset with young (<9 months) and old (>19 months) age bins; **(e)** publicly available single-cell RNAseq dataset of whole brain from male mice at young (2-3 months) and old (21-22 months) age, with the dataset sharing 264 genes in common with the coronal section dataset. Missing genes were imputed using SpaGE before spatial aging clock predictions were obtained. For statistics, see Supplementary Table 12.

**Extended Data Figure 7:**
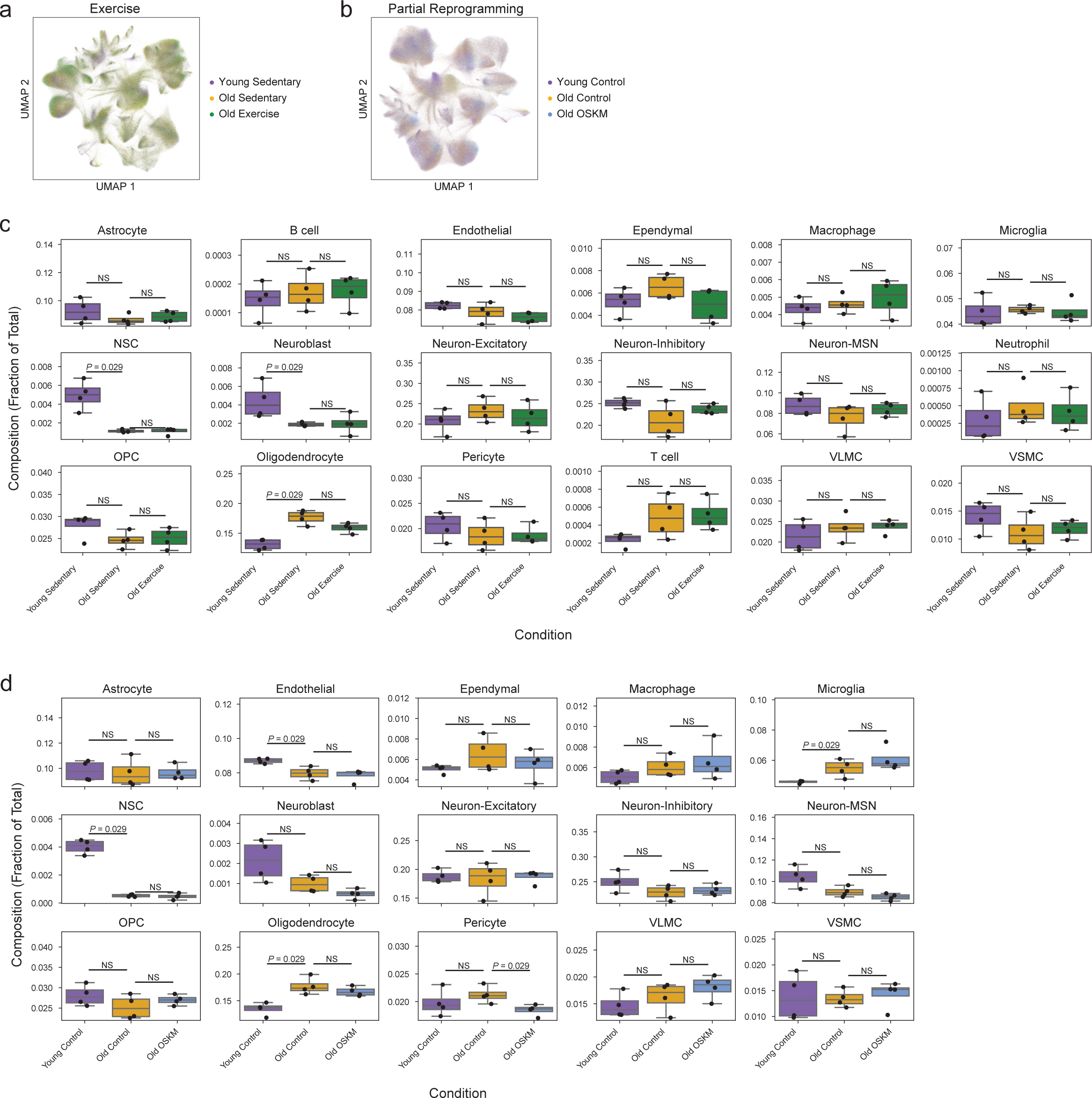
Spatial aging clocks record rejuvenation interventions. **a,b,** UMAP visualization of all cells by age and experimental condition for the **(a)** exercise cohort and **(b)** partial reprogramming cohort. **c,d,** Box plots showing cell type proportions for each cell type and across experimental conditions for the **(c)** exercise cohort and **(d)** partial reprogramming cohort. Dots indicate proportions for individual mice. Center denotes median, box spans the interquartile range, and whiskers span up to 1.5 times the interquartile range with diamonds showing outlier values. *P*-values computed using two-sided Mann-Whitney test. Not significant *P* > 0.05 abbreviated as “NS”.

**Extended Data Figure 8:**
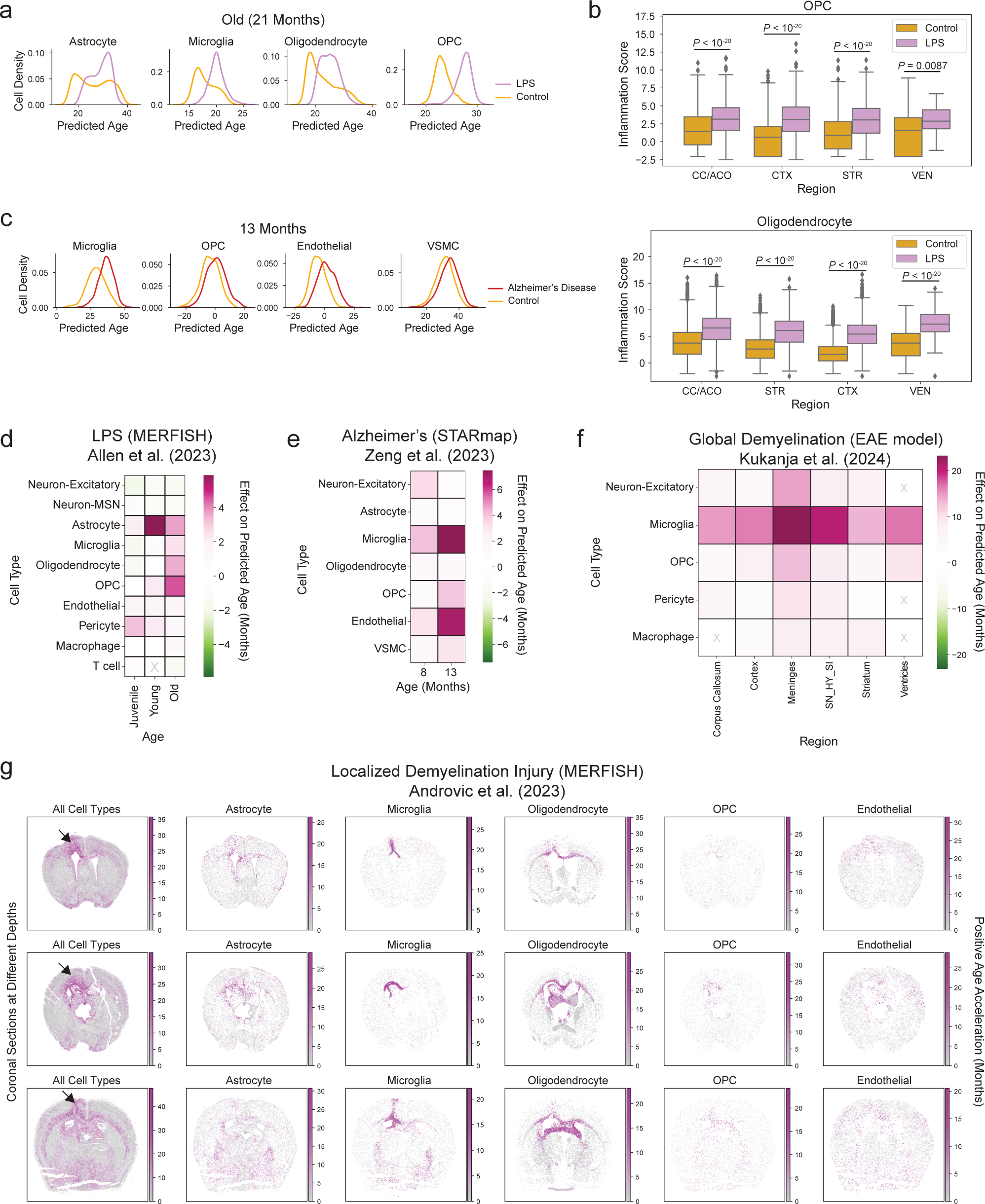
Spatial aging clocks record adverse interventions. **a,** Density of predicted ages across different experimental conditions and ages for spatial single-cell aging clocks corresponding to four cell types with accelerated aging under LPS-induced inflammation. For statistical analysis, see Supplementary Table 12. **b,** Boxplots of the oligodendrocyte inflammation score computed on scaled log-normalized gene expression values from the LPS MERFISH dataset for oligodendrocyte progenitor cells (OPCs) (top) and oligodendrocytes (bottom) across four anatomic regions and compared between LPS-injected and control conditions. *P*-values computed using two-sided Mann-Whitney test. **c,** Density of predicted ages across different experimental conditions and ages for spatial single-cell aging clocks corresponding to four cell types with accelerated aging in Alzheimer’s disease models. For statistical analysis, see Supplementary Table 12. **d,e,** Heatmaps showing the effect of adverse interventions on predicted age for different cell types, measured as the difference in median predicted age between intervention and control conditions, across different age groups, under **(d)** systemic inflammatory challenge by LPS injection across female juvenile (0.93 months), young (5.58 months), and old (20.93 months) mice and **(e)** for Alzheimer’s disease model male mice across 8 and 13 months of age. Gray “X” denotes cell types and regions with insufficient numbers of cells (<50). **f,** Heatmap showing the effect of global demyelination injury (EAE model of multiple sclerosis) on predicted age for different cell types and regions, measured as the difference in median predicted age between intervention and control conditions for the EAE model in young (2.2 months) male and female mice in an *in situ* sequencing dataset. Gray “X” denotes cell types and regions with insufficient numbers of cells (<50). “SN_HY_SI” refers to the region between the ventricles containing the substantia nigra and hypothalamus. **g,** Scatter plot of cells by their spatial coordinates in the localized demyelination injury MERFISH dataset with cells colored by the positive age acceleration (age acceleration with a floor value of zero) obtained from the spatial aging clocks. The leftmost column shows all cells with the site of injury marked with an arrow, and the other columns show select cell types. Three coronal sections at different depths are shown across the rows.

**Extended Data Figure 9:**
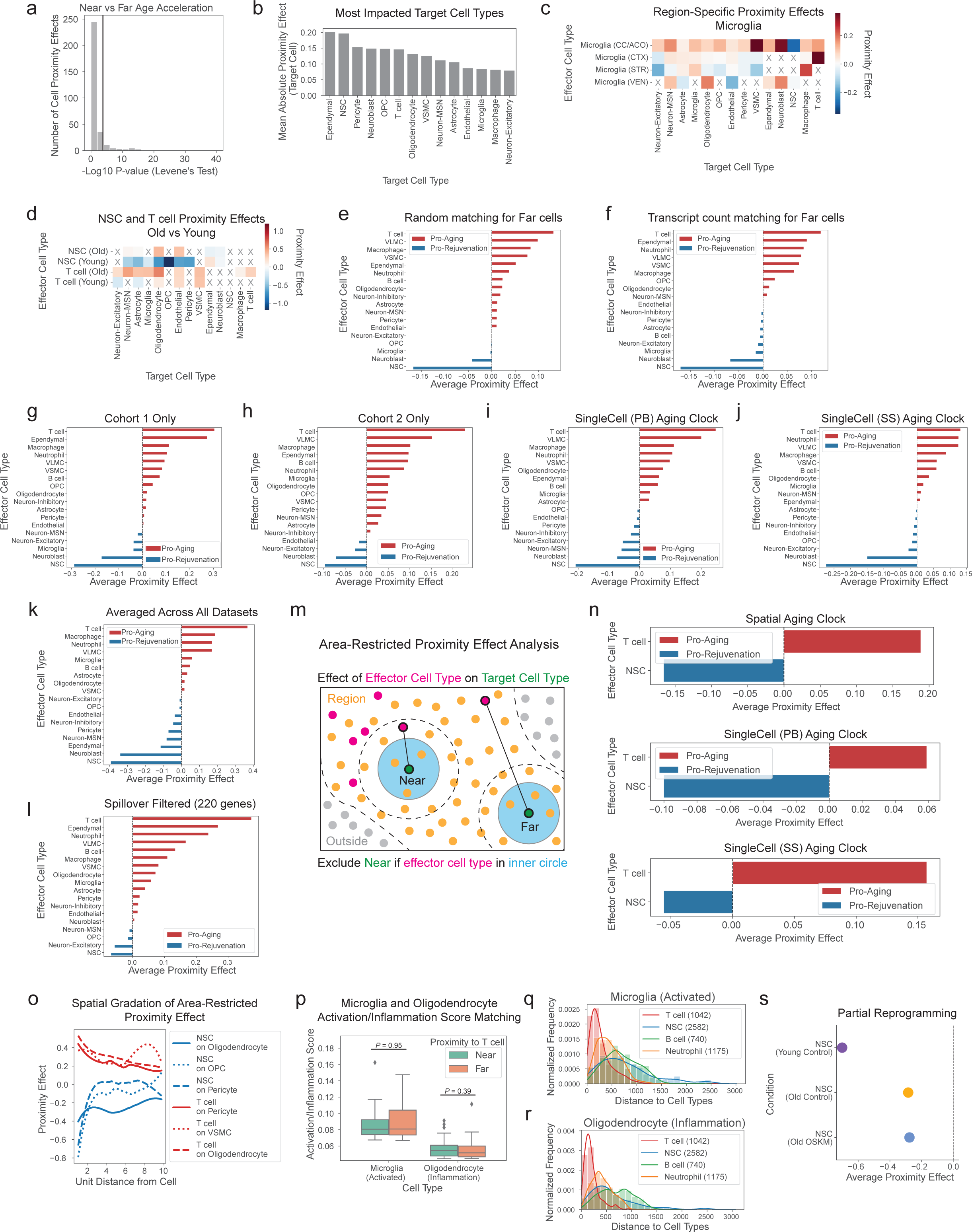
Specificity and robustness of cell proximity effects. **a,** Histogram showing distribution of negative log10 *P*-values from Levene’s test for equal variance between age acceleration of “Near” versus “Far” cell groups for each of the observed cell proximity effects. Solid vertical black line indicates the Bonferroni-adjusted cutoff corresponding *P* = 0.05. **b,** Target cell types ranked by their mean absolute proximity effect experienced across all effector cell types. **c,d,** Heatmap showing the proximity effect for different cell type proximity relationships **(c)** separated by anatomic region for microglia or **(d)** separated by young (<16 months) and old (>16 months) for NSCs and T cells. Rows correspond to the effector cell type and columns correspond to the target cell type, which experiences the proximity effect by the effector cells. “X” denotes proximity relationships for which there were insufficient cell pairings (<50) to compute a proximity effect (Cohen’s D of age acceleration, see Methods). **e-l,** Average proximity effect (APE) for a given effector cell type on all other target cell types ranked from most pro-aging (positive aging effect) to most pro-rejuvenating (negative aging effect) **(e)** with random matching to determine “Far” cells in the coronal section dataset; **(f)** with total transcript count matching to determine “Far” cells in the coronal section dataset; **(g)** with only mice in the first independent cohort (*n* = 10) of the coronal section dataset; **(h)** with only mice in the second independent cohort (*n* = 10) of the coronal section dataset; **(i)** using non-spatial aging clocks with pseudobulked single-cell gene expression for training and prediction (SingleCell (PB), see Methods); **(j)** using non-spatial aging clocks without SpatialSmooth for prediction (SingleCell (SS), see Methods); **(k)** for the mean APE across seven datasets (see Supplementary Table 15); and **(l)** using spatial aging clocks trained on a subset of 220 genes after filtering by transcript spillover rate (see Methods). **m,** Schematic illustration of the matching process for determining “Near” and “Far” target cells to compute the area-restricted proximity effect of effector cells on the age acceleration of target cells. **n,** Shown from top to bottom are the average area-restricted proximity effects (outer radius set to 2 unit distances with inner radius set at minus 15µm radius from the outer radius) for T cells and NSCs using spatial aging clocks (SpatialSmooth), using non-spatial aging clocks with pseudobulked single-cell training and prediction (SingleCell (PB), see Methods), and using non-spatial aging clocks without SpatialSmooth for prediction (SingleCell (SS), see Methods). For all three setups, T cells were among the most pro-aging cell types (ranked 1/18, 1/18, 4/18, respectively), and NSCs were among the most pro-rejuvenating cell types (ranked 1/18, 2/18, 1/18, respectively). **o,** Area-restricted proximity effect as a function of the outer radius distance (inner radius set at minus 15µm radius from the outer radius) for the top two pro-aging proximity relationships with T cell as the effector cell type and the top two pro-rejuvenating proximity relationships with NSC as the effector cell type. **p,** Boxplots of the glial activation/inflammation gene signature score for “Far” and “Near” activated/inflammation glia near T cells (activated/inflammation status determined by top 0.2% expression of signature). Center denotes median, box spans the interquartile range, and whiskers span up to 1.5 times the interquartile range with diamonds showing outlier values. *P*-values computed using two-sided Mann-Whitney test. **q,r,** Density plots and histograms showing the nearest distance to different cell types for **(q)** all activated microglia and **(r)** all inflammation oligodendrocytes in the coronal section dataset. Number of cells for each cell type are listed in parentheses in the legend. **s,** Average proximity effect of NSCs on nearby cells computed from all cells in the partial reprogramming dataset for each of the three experimental conditions (Young Control, Old Control, Old OSKM). There were no T cells detected in the partial reprogramming mouse model (see Methods).

**Extended Data Figure 10:**
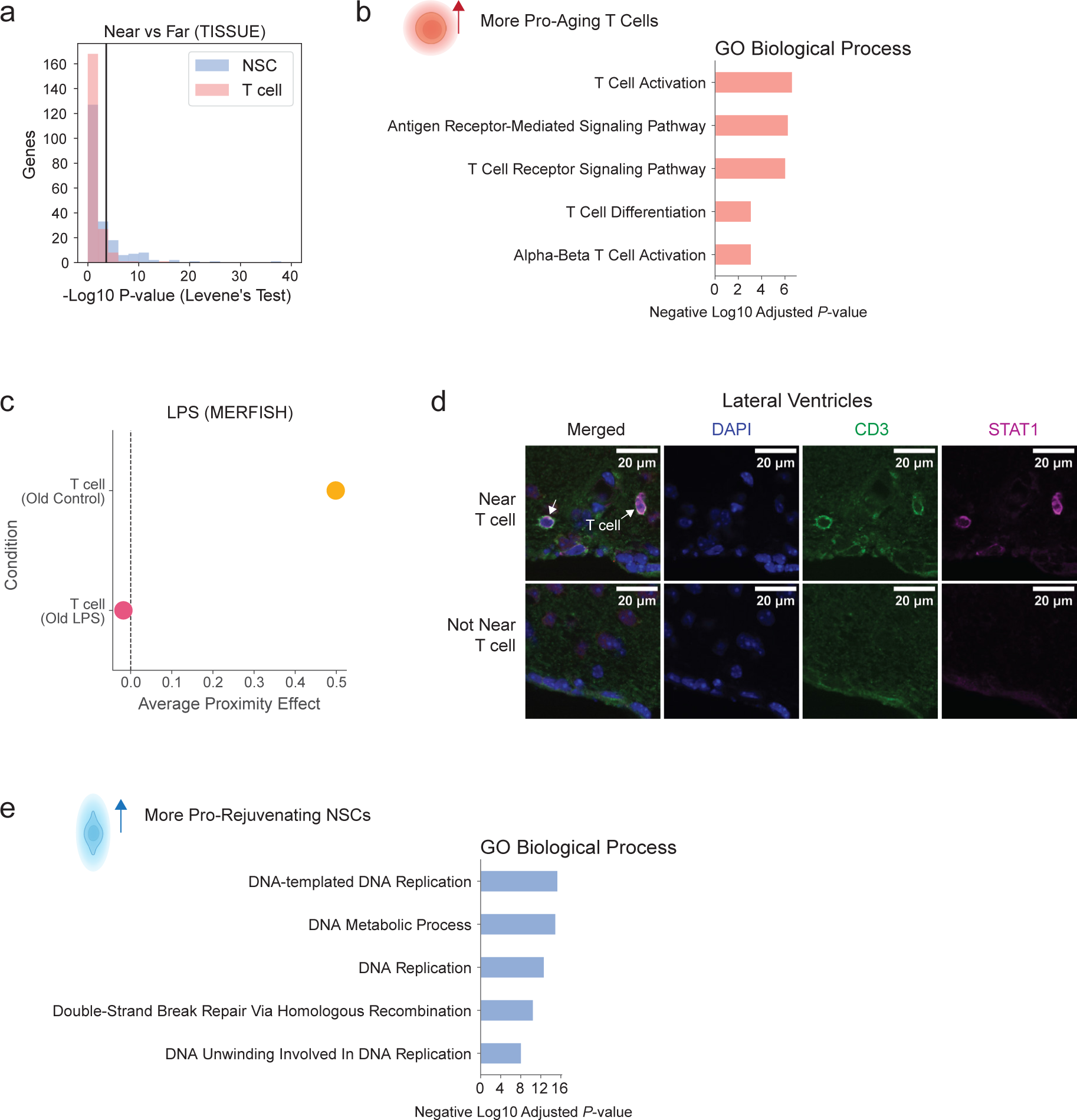
Identification and validation of potential mediators of cell proximity effects. **a,** Histogram showing distribution of negative log10 *P*-values from Levene’s test for equal variance between SpaGE-imputed gene expression in “Near” versus “Far” cells with respect to NSCs or T cells. Solid vertical black line indicates the Bonferroni-adjusted cutoff corresponding *P* = 0.05. **b,** Bar plots showing the top five most enriched GO Biological Processes for significantly upregulated genes in more pro-aging T cells compared to less pro-aging T cells. *P*-values computed from EnrichR pathway enrichment analysis. **c,** Average proximity effects of T cells computed from all nearby cells in the LPS MERFISH study for each of the two experimental conditions (Old Control, Old LPS) with reduced *Ifng* expression in Old LPS condition. **d,** Immunofluorescence staining image of a perfused mouse brain section from an old (28 months old) male mouse highlighting elevated STAT1 fluorescence (magenta) in cells that are near T cells (marked by arrows, top panels) compared to cells that are not near T cells (bottom panels) in the lateral ventricles. Images were taken from the same brain section. **e,** Bar plots showing the top five most enriched GO Biological Processes for significantly upregulated genes in more pro-rejuvenating NSCs compared to less pro-rejuvenating NSCs. *P*-values computed from EnrichR pathway enrichment analysis.

**Supplementary Figure 1:**
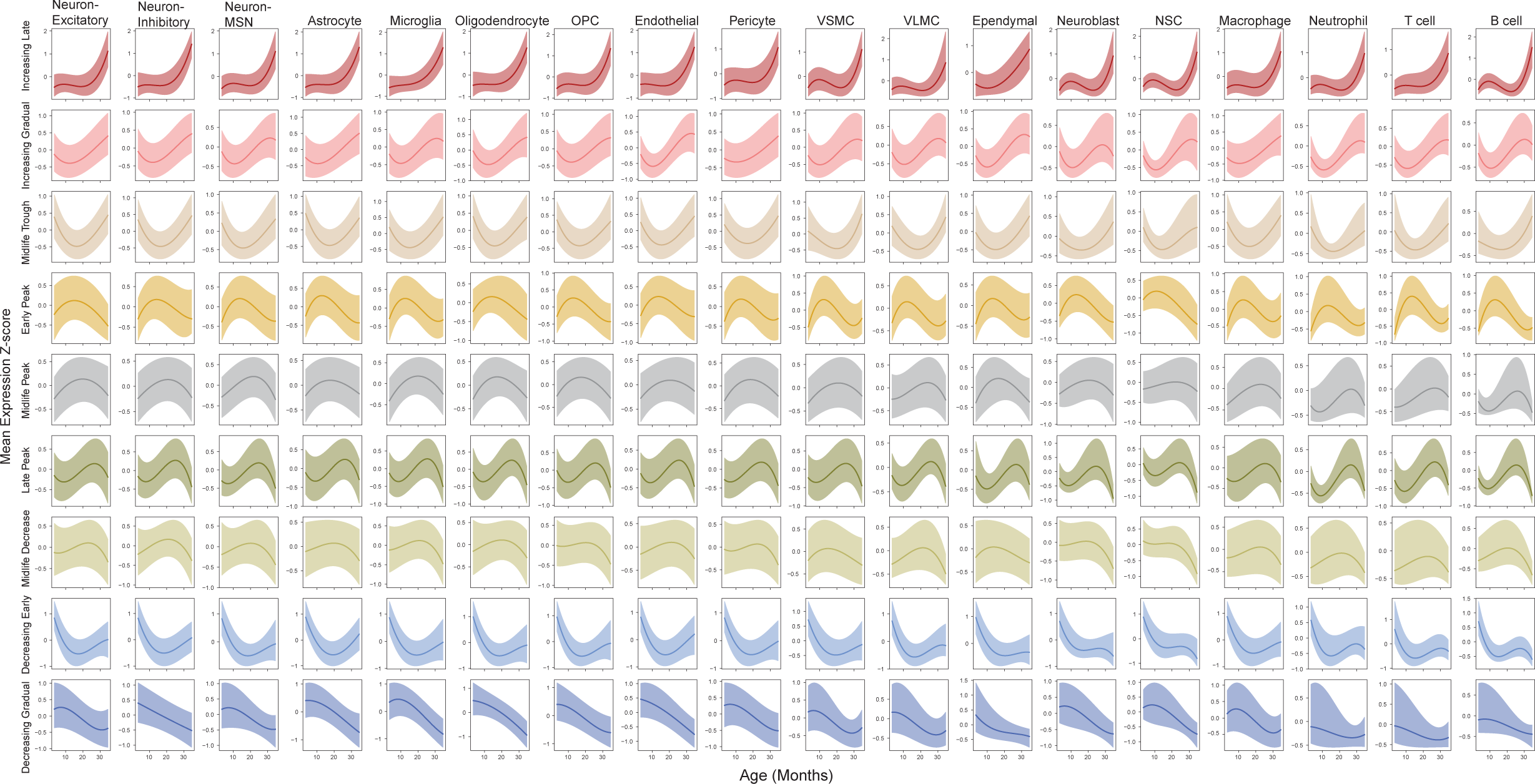
Median gene expression z-score (see Methods) across age of all gene, region, cell type combinations (coronal sections) split into the nine annotated gene trajectory clusters (rows) and further divided by cell type (columns). Shaded regions correspond to interquartile range in expression. All trajectories were smoothed using interpolating B-splines.

**Supplementary Figure 2:**
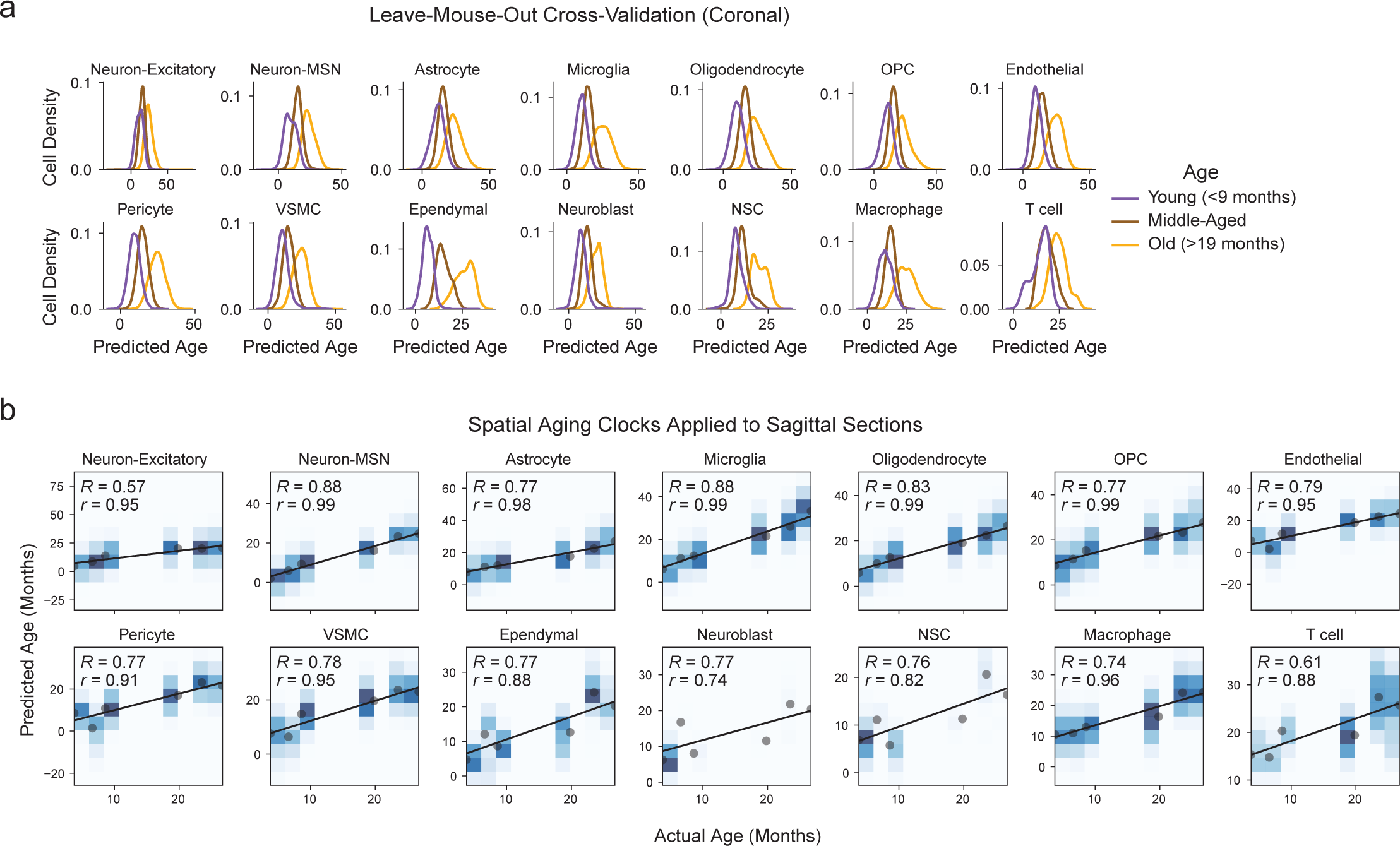
**a**, Density of predicted ages computed using spatial aging clocks for different age groups in the coronal section dataset using cross-validation with groups defined by binning samples into young (less than 9 months), middle-aged (between 9 and 19 months), and old (greater than 19 months) age groups. Missing genes were imputed using SpaGE before spatial aging clock predictions were obtained. For statistical analysis, refer to Supplementary Table 12. **b**, Predicted age as a function of actual age for the sagittal section dataset using the spatial aging clocks. Heatmap colors represent density of predicted ages. Gray circles represent the median predicted age for an individual mouse. The Pearson correlation between predicted age and actual age for all cells is reported as R, and the Pearson correlation between median predicted age and actual age for all mice is reported as r.

**Supplementary Figure 3:**
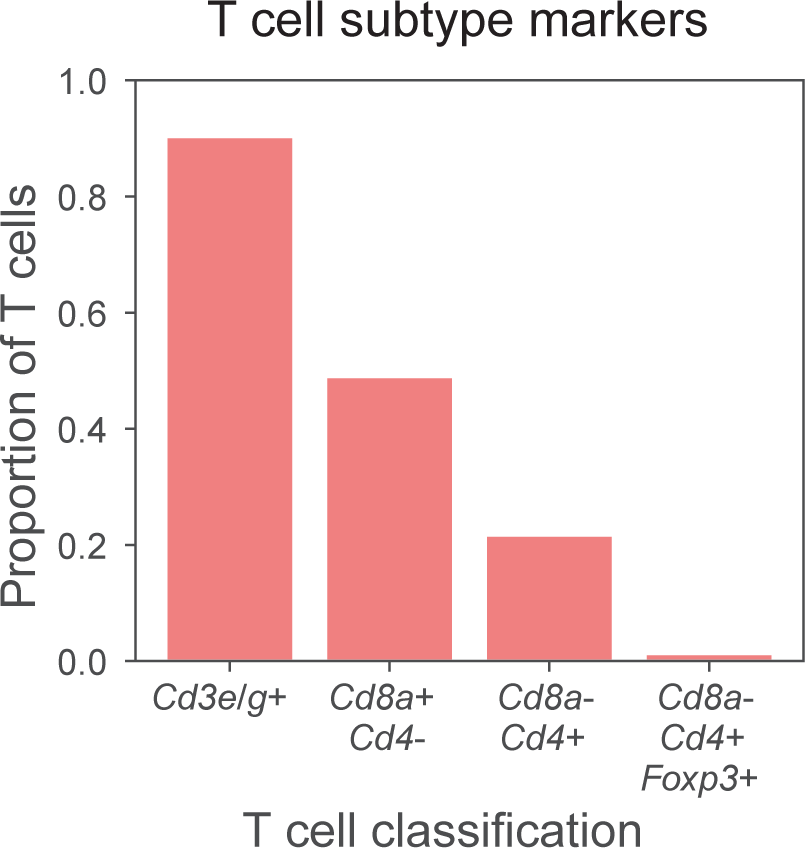
Bar plots showing the proportion of T cells split into different subtypes defined by presence of T cell subtype marker transcripts in the MERFISH coronal section dataset.

